# High-accuracy HLA type inference from whole-genome sequencing data

**DOI:** 10.1101/035253

**Authors:** Alexander T Dilthey, Pierre-Antoine Gourraud, Zamin Iqbal, Gil McVean

## Abstract

Extensive hyperpolymorphism and sequence similarity between the HLA genes make HLA type inference from whole-genome sequencing data a challenging problem. We address these by representing sequences from over 10,000 known alleles in a reference graph structure, enabling accurate read mapping. HLA*PRG, our algorithm, outperforms existing methods by a wide margin and for the first time consistently achieves the accuracy of gold-standard reference methods with one error across 158 alleles tested.

---

Genetic variation at HLA loci, both classical and non-classical, is associated with many medical phenotypes including risk of autoimmune^1–3^ and infectious^4^ disease, adverse drug reactions^5, 6^, success of tissue and organ transplants^7^, and, via epitope presentation preferences, the success of cancer immunotherapy^8^. The current gold standard for high resolution typing of HLA alleles, sequence-based typing (SBT), uses Sanger sequencing or targeted amplification of the HLA genes followed by next-generation sequencing and can require extensive manual curation, thus making high throughput application of the method expensive and challenging. With the growth of high throughput genomic technologies, methods for inferring HLA genotype have been developed that use SNP genotyping^9–12^, exome and whole-genome sequencing^13–17^. These approaches offer high throughput, but, to date, are either limited to a subset of HLA loci^17^ or do not achieve the same degree of accuracy as SBT. Multiple factors influence accuracy, including the sheer sequence and structural diversity of the region, the presence of multiple paralogous genes (including pseudogenes) and rare, but important, gene conversion events that generate mosaic allelic structures.

To address these challenges, we have previously introduced structures to represent known genomic variation called population reference graphs (PRGs) and demonstrated their value in characterising variation across the MHC and particularly within the HLA Class II gene region^18^. Briefly, a PRG is a directed graph in which alternative alleles, insertions and deletions are represented as alternative paths through the graph, and in which orthologous and identical regions are collapsed locally to model potential recombination. Although expensive computationally, reads likely to arise from the region can be identified and mapped directly to the graph structure, thus enabling the assessment of evidence for the presence of each stretch of sequence along a path. The pairs of paths with greatest joint support can therefore be identified and assigned as the diploid genotype for an individual. Previously, we demonstrated that a prototype of this approach can identify the nucleotide-level variants at classical HLA alleles with high accuracy. However, we did not address the problem of inferring the allele present at the gene level^18^.

Allelic variants at HLA genes can be typed^19, 20^ at different degrees of resolution; low resolution (“2-digit”) types specify serological activity; intermediate resolution (“4-digit”) HLA types specify the complete primary sequence of the HLA proteins and high-resolution (“6-digit”) types determine the full exonic sequence including synonymous variants. Higher levels of resolution include non-coding variation. SBT is typically carried out at 6-digit “G” resolution, in which only the sequences of the exons encoding the peptide binding groove are considered: exons 2 and 3 for HLA class I genes and exon 2 for HLA class II genes. In most applications of typing, a set of 6–8 loci are typed (Class I: *HLA-A, -B*, -C, Class II: *HLA-DQA1, -DQB1, -DRB1, -DRA1* and *-DPB1)*, though there exist over 30 HLA genes and pseudogenes. Supplementary Figures 1 and 2 demonstrate the high degree of sequence similarity and its non-random spatial structure between alleles in certain groups of loci.

We set out to modify the existing PRG approach to provide accurate HLA typing at 6-digit “G” resolution using high coverage whole-genome sequencing data, such as is being generated by large-scale genomics projects. Details of the approach, including source data, graph-construction, read-mapping and HLA typing are given in the Supplementary Note and a schematic is shown in Figure. 1. Briefly, we build a gene-specific PRG comprising 46 (mostly HLA) genes and pseudogenes, 720 genomic and 10,500 coding (exon-only) alleles from IMGT/HLA^21^ (Supplementary Table 1). Each gene is embedded in a stretch of surrounding reference sequence, but we don’t attempt to model the full intergenic sequence. Reads are mapped to the PRG and the pair of alleles with the highest joint likelihood is identified and reported with an associated quality measure for each individual allele (integrating over the distribution of posterior probabilities, see Supplementary Note). The software to carry out these steps, HLA*PRG, is freely available.

**Figure 1.**
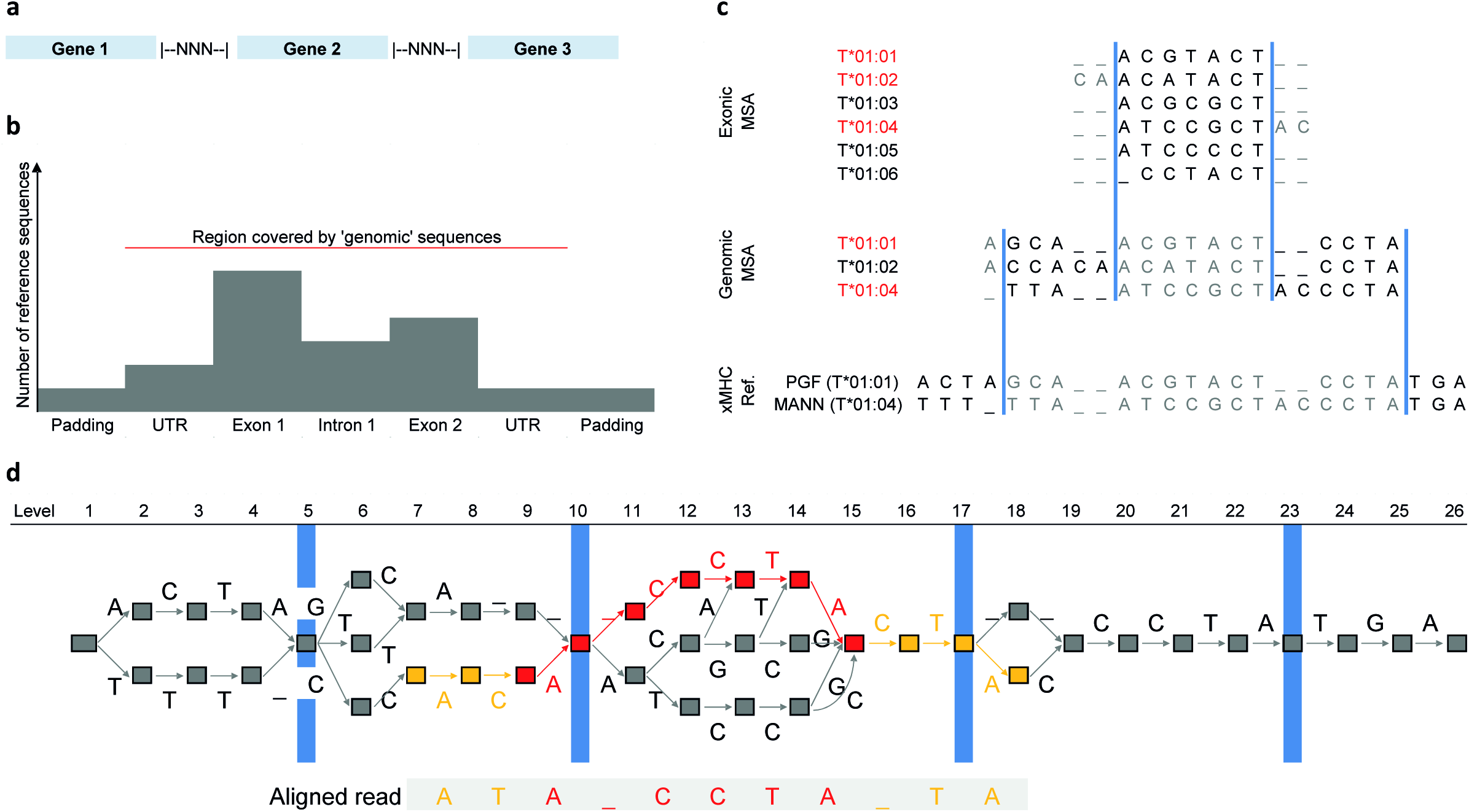
Schematic representation of HLA type inference using HLA*PRG. **a** Broad-scale structure of the HLA PRG. The included genes are separated by spacer blocks consisting of N characters. **b** Fine-scale structure of the PRG input sequences. Exons, introns and UTRs are embedded in regional haplotypes (padding sequence). Exon sequences typically outnumber intron sequences. The red line indicates the region covered by IMGT genomic sequences. **c** For each gene represented in the PRG, multiple sequence alignments representing up to 3 sources of sequence data are merged for PRG construction: exonic sequences, genomic (UTR, exons, introns) sequences, regional haplotypes (“xMHC Ref.”). Using alleles present in both the current and the next-higher-level MSA (identifiers printed in red), the merging algorithm determines consensus boundaries (blue bars) to connect the MSAs of different input sequence types. For each segment so-defined, we use the MSA corresponding to the highest-resolution input sequence type (sequence characters therefore ignored are printed in grey). **d** The PRG corresponding to the input sequences shown in c, and a seed-and-extend alignment of a sequencing read to the PRG. PRG nodes are represented by boxes and edges by labelled arrows. The four blue markers correspond to the consensus MSA boundaries shown in c. The aligned sequence of the read is displayed below the PRG, and the alignment path (the sequence of edges and nodes traversed in the PRG) is highlighted. The red component of the alignment path corresponds to the exact-match “seed” component of the alignment (spanning a graph-encoded gap), whereas the orange components correspond to the “extend” component of the alignment (where mismatches are allowed).

**Table 1.**

| Cohort | Locus | N | HLA*PRG |  |  | PHLAT |  |  | HLAreporter |  |  |
| --- | --- | --- | --- | --- | --- | --- | --- | --- | --- | --- | --- |
|  |  |  | Inferred | Accuracy | Call Rate | Inferred | Accuracy | Call Rate | Inferred | Accuracy | Call Rate |
| Platinum Trio | A | 6 | 6 | 1.00 | 1.00 | 6 | 1.00 | 1.00 | 2 | 0.50 | 0.33 |
|  | B | 6 | 6 | 1.00 | 1.00 | 6 | 1.00 | 1.00 | 1 | 1.00 | 0.17 |
|  | C | 6 | 6 | 1.00 | 1.00 | 6 | 1.00 | 1.00 | 1 | 0.00 | 0.17 |
|  | DQA1 | 6 | 6 | 1.00 | 1.00 | 6 | 1.00 | 1.00 | 2 | 1.00 | 0.33 |
|  | DQB1 | 6 | 6 | 1.00 | 1.00 | 6 | 1.00 | 1.00 | 5 | 1.00 | 0.83 |
|  | DRB1 | 6 | 6 | 1.00 | 1.00 | 6 | 1.00 | 1.00 | 5 | 1.00 | 0.83 |
| 1000 Genomes Highest Resolution | A | 22 | 22 | 1.00 | 1.00 | 20 | 0.45 | 0.91 | 0 | NA | 0.00 |
|  | B | 22 | 22 | 1.00 | 1.00 | 20 | 0.35 | 0.91 | 6 | 0.50 | 0.27 |
|  | C | 22 | 22 | 1.00 | 1.00 | 20 | 0.50 | 0.91 | 2 | 0.50 | 0.09 |
|  | DQA1 | 12 | 12 | 1.00 | 1.00 | 10 | 0.70 | 0.83 | 9 | 1.00 | 0.75 |
|  | DQB1 | 22 | 22 | 1.00 | 1.00 | 20 | 0.80 | 0.91 | 15 | 1.00 | 0.68 |
|  | DRB1 | 22 | 22 | 0.95 | 1.00 | 20 | 0.55 | 0.91 | 10 | 1.00 | 0.45 |
| 1000 Genomes 4-digit G<sup>a</sup> | A | 22 | 22 | 1.00 | 1.00 | 20 | 0.70 | 0.91 | 0 | NA | 0.00 |
|  | B | 22 | 22 | 1.00 | 1.00 | 20 | 0.60 | 0.91 | 6 | 0.50 | 0.27 |
|  | C | 22 | 22 | 1.00 | 1.00 | 20 | 0.80 | 0.91 | 2 | 0.50 | 0.09 |
|  | DQA1 | 12 | 12 | 1.00 | 1.00 | 10 | 0.70 | 0.83 | 9 | 1.00 | 0.75 |
|  | DQB1 | 22 | 22 | 1.00 | 1.00 | 20 | 0.95 | 0.91 | 15 | 1.00 | 0.68 |
|  | DRB1 | 22 | 22 | 0.95 | 1.00 | 20 | 0.75 | 0.91 | 10 | 1.00 | 0.45 |

HLA type inference accuracy for HLA\*PRG and two state-of-the-art algorithms
<sup>a</sup> “1000 Genomes Highest Resolution” and “1000 Genomes 4-digit G” represent the same set of samples, with 6-digit validation alleles (where available) reduced to 4-digit resolution for the latter experiment, enabling a fair comparison with algorithms that fall back to 4-digit typing in cases of ambiguity.

To assess the accuracy of the method, we used two data sets with available high coverage sequencing data and independent SBT-based HLA type information (Table 1). First, we analysed NA12878, NA12891 and NA12892 from the Illumina Platinum Genomes Project, sequenced to 50 – 55x with a PCR-free 2 × 100bp protocol. We correctly infer all 36 HLA alleles. Second, we analysed 11 samples from the 1000 Genomes Project, sequenced to 28 – 68x with a PCR-free 2 × 250bp protocol. Initial analysis identified three discrepancies (Supplementary Note), though on re-typing these individuals two of three were the result of initial errors in the validation data. The remaining inconsistency, *(HLA-DRB1*16:02:01* incorrectly typed as *HLA-DRB1*16:23)* is likely caused by *HLA-DRB5* sequences incorrectly aligned to *HLA-DRB1* within IMGT (IMGT/HLA currently don’t provide genomic sequences for HLA-DRB5 and the representation of this gene in the PRG therefore remains incomplete).

We compare the performance of HLA*PRG with PHLAT^14^ and HLAreporter^13^, two state-of-the-art algorithms that support HLA class I and class II. For the Platinum samples, we find that PHLAT also correctly infers all 36 alleles, whereas HLAreporter only reports 16 alleles (of which 14 are correct). For the 1000 Genomes Samples, we find that HLA*PRG outperforms both programs by a wide margin. Mean accuracy at 4-digit resolution across all loci is 75% for PHLAT and 80% for HLAreporter, and HLAreporter achieves a call rate of only 38%. To assess to what extent HLA*PRG depends on the availability of whole-genome data, we also apply it to whole-exome sequencing data of a cohort of HapMap samples. Results are varied and accuracies consistently lower across all loci (ranging from 79% for *HLA-C* to 98% for *HLA-DQB1*, Supplementary Table 2). To assess sensitivity of HLA*PRG to whole-genome sequencing depth, we subsampled the NA12878 data from the Platinum and 1000 Genomes projects to average coverages of 40x, 30x and 20x in triplicates. We find that performance is stable (all alleles correctly predicted) down to 20x for the Platinum data and down to 30x for the 1000 Genomes data (Supplementary Table 3). To assess whether HLA*PRG could be applied to additional HLA loci beyond the set of 6 genes validated here, we used it to genotype a set of 12 additional HLA genes and pseudogenes in the Illumina Platinum data (Supplementary Table 4). Across the 72 alleles inferred, we find one trio inconsistency at the pseudogene *HLA-K*, which is driven by an allele called with low confidence.

Our current implementation is optimised for accuracy rather than computational efficiency. Analysing the NA12878 Platinum data (55x depth) takes 11 hours (clock time, AMD Opteron 6174 2.2GHz), and 17 hours for the NA12878 1000 Genomes data (63x coverage). We provide a detailed runtime (including CPU time) and memory analysis in the Supplement (Supplementary Note). Achieving improvements in computational efficiency is ongoing work. Future versions might make use of linear sequence alignments to seed graph alignment and also leverage population haplotype frequencies^22, 23^.

In conclusion, we find that HLA*PRG infers HLA types at accuracies comparable to current gold standard typing technologies (two errors in the original reference data compared to one from HLA*PRG at 4-digit / 6-digit resolution), provided that high-quality (PCR-free protocol, read length of at least 100bp, coverage of at least 30x) whole-genome sequencing data are used as input. HLA*PRG will enable researchers to augment population-scale whole-genome sequencing data with reliable HLA type information and contribute to characterizing HLA signals in important medical phenotypes.

## Acknowledgements

The study was funded by grant 100956/Z/13/Z from the Wellcome Trust to G.M., a Nuffield Department of Medicine Fellowship to Z.I. and a Sir Henry Dale Fellowship jointly awarded by the Wellcome Trust and the Royal Society to Z.I. (102541/Z/13/Z). P.A.G. is supported by ATIP-Avenir INSERM program and the Region Pays de Loire ConnecTalent.

## Data availability

All sequencing data used in the study are publically available through existing projects. URLs for sequencing data access and utilized HLA types are given in Section 4 of the Supplementary Note.

## Software availability

HLA*PRG is implemented in C++/Perl and available under GPLv3 as part of the MHC*PRG repository https://github.com/AlexanderDilthey/MHC-PRG. A readme file (https://github.com/AlexanderDilthey/MHC-PRG/blob/master/HLA-PRG.md) describes how to install and run the software.

---

**Supplementary Figure 1.**
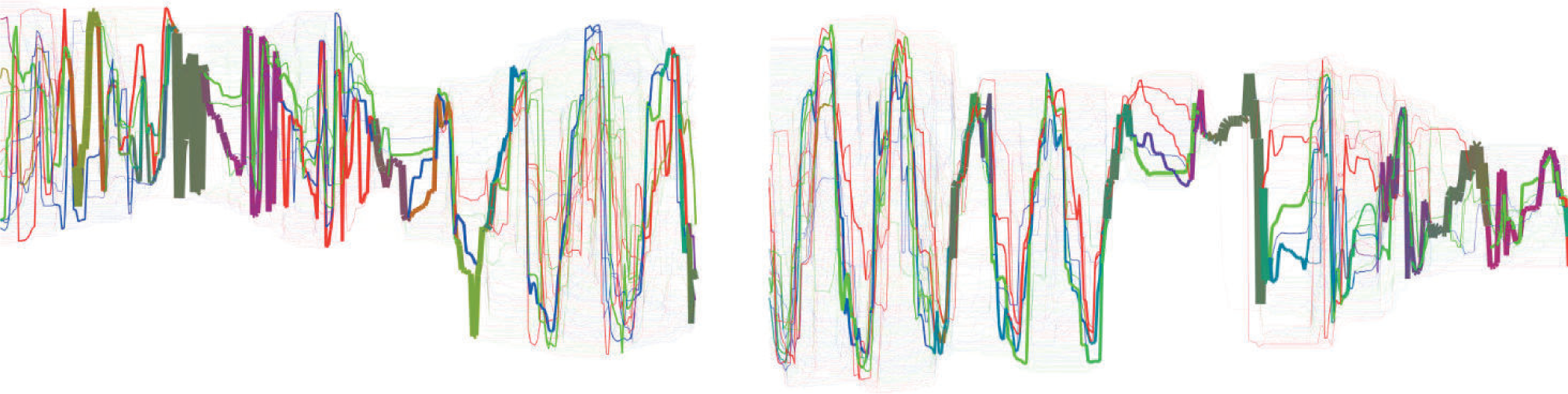
Sequence homology between HLA-A, -B and -C. Graph visualizing sequence homology between *HLA-A, -B* and *-C* across exons 2 (left) and 3 (right), based on a multiple sequence alignment (MSA) of 3284 -A, 4077 -B, 2799 -C alleles. The x axis of the plot represents the column index of the MSA (304 columns for exon 2, 349 columns for exon 3). The (invisible) nodes of the graph represent the set of unique 31-mers (across the 3 genes) starting at the corresponding column of the MSA. Two nodes (representing two consecutive 31-mers in the MSA) are connected by (visible) edges if the corresponding 32-mer, starting at the column index of the first 31-mer, is present in the MSA. Edge flow (line thickness) is proportional to the frequency of the corresponding 32-mer at the underlying column (bounded below). Edge colour indicates the proportions of flow attributable to the 3 genes (for each edge, the absolute count of the corresponding 32-mer at the underlying column can be split into a triplet representing the *HLA-A, HLA-B, HLA-C* rows of the alignment; the (R, G, B) colour of the edge is obtained by normalizing this triplet). For the purpose of this plot, we treat gap characters as nucleotides.

**Supplementary Figure 2.**
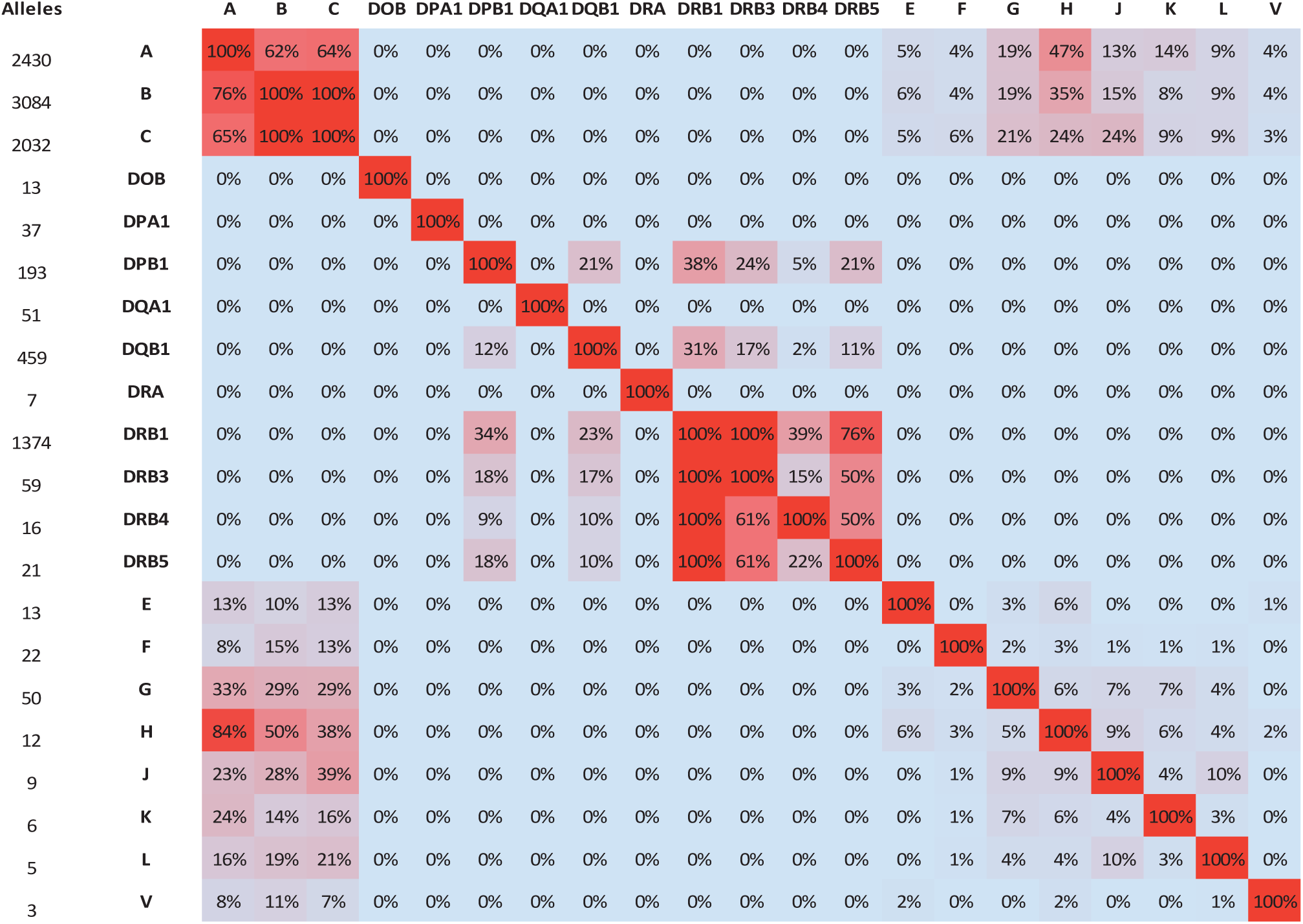
Maximum HLA gene sequence homology at the peptide binding site. Maximum k-Mer similarity at the peptide binding site (PBS; exons 2/3 for HLA class I, exon 2 for HLA class II) between alleles of different HLA loci, based on k-Mers (k = 25). 6-digit HLA “G” types are defined by PBS sequence. Each cell, in row X and column Y, contains the maximum, over all alleles of locus X, proportion of k-Mers present in any allele of locus Y. This quantity influences the probability of mismapping a PBS read to another locus as exact matching is the first step of the many mapping algorithms, including the one used here.

**Supplementary Table 1.**
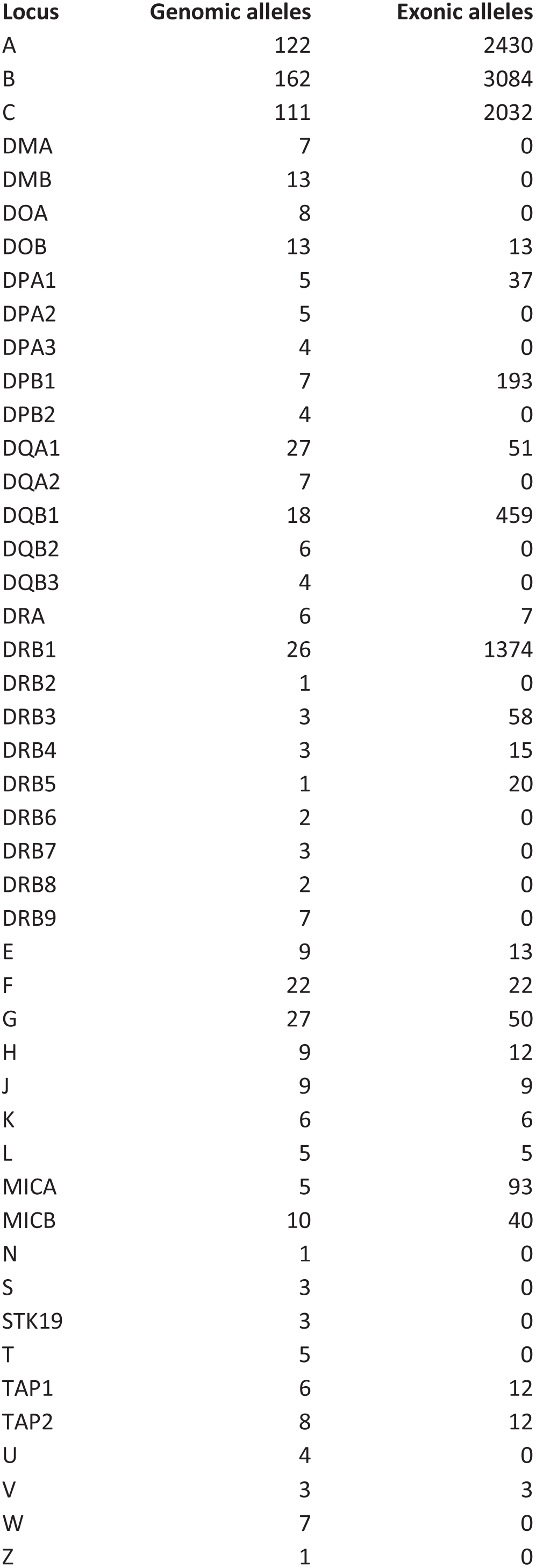
HLA PRG input sequences Loci represented in the HLA PRG. “Genomic alleles”: Genomic alleles represented in the gene-specific segment of the PRG, i.e. alleles spanning the complete length of the gene. “Exonic alleles”: Exonic alleles represented in the gene-specific fragment of the PRG.

| Locus | Genomic alleles | Exonic alleles |
| --- | --- | --- |
| A | 122 | 2430 |
| B | 162 | 3084 |
| C | 111 | 2032 |
| DMA | 7 | 0 |
| DMB | 13 | 0 |
| DOA | 8 | 0 |
| DOB | 13 | 13 |
| DPA1 | 5 | 37 |
| DPA2 | 5 | 0 |
| DPA3 | 4 | 0 |
| DPB1 | 7 | 193 |
| DPB2 | 4 | 0 |
| DQA1 | 27 | 51 |
| DQA2 | 7 | 0 |
| DQB1 | 18 | 459 |
| DQB2 | 6 | 0 |
| DQB3 | 4 | 0 |
| DRA | 6 | 7 |
| DRB1 | 26 | 1374 |
| DRB2 | 1 | 0 |
| DRB3 | 3 | 58 |
| DRB4 | 3 | 15 |
| DRB5 | 1 | 20 |
| DRB6 | 2 | 0 |
| DRB7 | 3 | 0 |
| DRB8 | 2 | 0 |
| DRB9 | 7 | 0 |
| E | 9 | 13 |
| F | 22 | 22 |
| G | 27 | 50 |
| H | 9 | 12 |
| J | 9 | 9 |
| K | 6 | 6 |
| L | 5 | 5 |
| MICA | 5 | 93 |
| MICB | 10 | 40 |
| N | 1 | 0 |
| S | 3 | 0 |
| STK19 | 3 | 0 |
| T | 5 | 0 |
| TAP1 | 6 | 12 |
| TAP2 | 8 | 12 |
| U | 4 | 0 |
| V | 3 | 3 |
| W | 7 | 0 |
| Z | 1 | 0 |

**Supplementary Table 2.** Performance on exome sequencing data HLA type inference accuracy, per locus, for HLA*PRG and two state-of-the-art algorithms, PHLAT and HLAreporter, on a set of exome-sequenced HapMap samples (2 × 100bp, average per-locus coverage at the peptide-binding site 54x (over all validated HLA loci and samples, minimum 4.4x, maximum 164x). “Highest Resolution” and “4-digit G” represent the same set of samples, with 6-digit validation alleles (where available) reduced to 4-digit resolution for the latter experiment. Note that the number of inferred alleles varies between algorithms.

| Cohort | Locus | N | HLA*PRG |  |  | PHLAT |  |  | HLAreporter |  |  |
| --- | --- | --- | --- | --- | --- | --- | --- | --- | --- | --- | --- |
|  |  |  | Inferred alleles | Accuracy | Call Rate | Inferred alleles | Accuracy | Call Rate | Inferred alleles | Accuracy | Call Rate |
| HapMap<br>Exomes<br>Highest<br>Resolution | A | 58 | 58 | 0.86 | 1.00 | 58 | 0.78 | 1.00 | 20 | 0.95 | 0.34 |
|  | B | 48 | 48 | 0.85 | 1.00 | 48 | 0.90 | 1.00 | 22 | 0.64 | 0.46 |
|  | C | 56 | 56 | 0.79 | 1.00 | 56 | 0.77 | 1.00 | 17 | 0.59 | 0.30 |
|  | DQA1 | 58 | 58 | 0.95 | 1.00 | 58 | 0.84 | 1.00 | 42 | 0.98 | 0.72 |
|  | DQB1 | 58 | 58 | 0.98 | 1.00 | 58 | 0.95 | 1.00 | 44 | 1.00 | 0.76 |
|  | DRB1 | 58 | 58 | 0.86 | 1.00 | 58 | 0.83 | 1.00 | 41 | 0.95 | 0.71 |
| HapMap<br>Exomes<br>4-digit G | A | 58 | 58 | 0.86 | 1.00 | 58 | 0.78 | 1.00 | 20 | 0.95 | 0.34 |
|  | B | 48 | 48 | 0.85 | 1.00 | 48 | 0.92 | 1.00 | 22 | 0.64 | 0.46 |
|  | C | 56 | 56 | 0.79 | 1.00 | 56 | 0.93 | 1.00 | 17 | 0.59 | 0.30 |
|  | DQA1 | 58 | 58 | 0.95 | 1.00 | 58 | 0.84 | 1.00 | 42 | 0.98 | 0.72 |
|  | DQB1 | 58 | 58 | 0.98 | 1.00 | 58 | 0.95 | 1.00 | 44 | 1.00 | 0.76 |
|  | DRB1 | 58 | 58 | 0.86 | 1.00 | 58 | 0.83 | 1.00 | 41 | 0.95 | 0.71 |

**Supplementary Table 3.** Coverage sensitivity analysis Sensitivity to reduced coverage. Results for NA12878 (Platinum and 1000 Genomes data, see main text), down-sampled to 40x, 30x, 20x (triplicates).

| Sample | Locus | N | HLA*PRG |  |
| --- | --- | --- | --- | --- |
|  |  |  | Correctly<br>inferred | Accuracy |
| NA12878<br>Platinum<br>C = 40 | A | 6 | 6 | 1.00 |
|  | B | 6 | 6 | 1.00 |
|  | C | 6 | 6 | 1.00 |
|  | DQA1 | 6 | 6 | 1.00 |
|  | DQB1 | 6 | 6 | 1.00 |
|  | DRB1 | 6 | 6 | 1.00 |
| NA12878<br>Platinum<br>C = 30 | A | 6 | 5 | 0.83 |
|  | B | 6 | 6 | 1.00 |
|  | C | 6 | 6 | 1.00 |
|  | DQA1 | 6 | 6 | 1.00 |
|  | DQB1 | 6 | 6 | 1.00 |
|  | DRB1 | 6 | 6 | 1.00 |
| NA12878<br>Platinum<br>C = 20 | A | 6 | 6 | 1.00 |
|  | B | 6 | 6 | 1.00 |
|  | C | 6 | 6 | 1.00 |
|  | DQA1 | 6 | 6 | 1.00 |
|  | DQB1 | 6 | 6 | 1.00 |
|  | DRB1 | 6 | 6 | 1.00 |
| NA12878<br>1000G<br>C = 40 | A | 6 | 6 | 1.00 |
|  | B | 6 | 6 | 1.00 |
|  | C | 6 | 6 | 1.00 |
|  | DQA1 | 6 | 6 | 1.00 |
|  | DQB1 | 6 | 6 | 1.00 |
|  | DRB1 | 6 | 6 | 1.00 |
| NA12878<br>1000G<br>C = 30 | A | 6 | 6 | 1.00 |
|  | B | 6 | 6 | 1.00 |
|  | C | 6 | 6 | 1.00 |
|  | DQA1 | 6 | 6 | 1.00 |
|  | DQB1 | 6 | 6 | 1.00 |
|  | DRB1 | 6 | 6 | 1.00 |
| NA12878<br>1000G<br>C = 20 | A | 6 | 6 | 1.00 |
|  | B | 6 | 6 | 1.00 |
|  | C | 6 | 6 | 1.00 |
|  | DQA1 | 6 | 6 | 1.00 |
|  | DQB1 | 6 | 6 | 1.00 |
|  | DRB1 | 6 | 3 | 0.50 |

**Supplementary Table 4.**
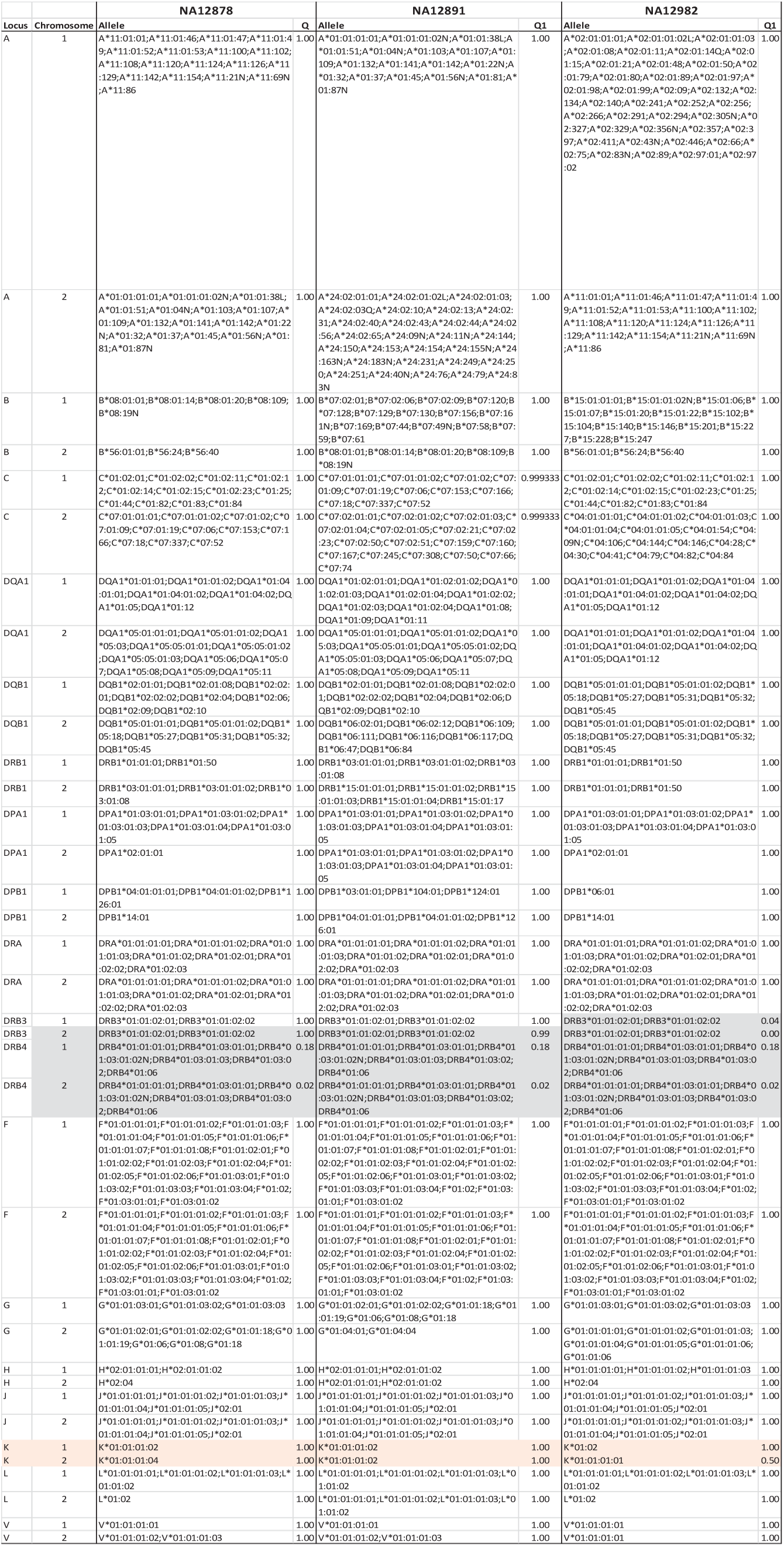
Inferred non-classical HLA type for the Platinum trio HLA types for the NA12878 (child), NA12891 (parent), NA12892 (parent) Platinum trio, including additional loci and typing quality scores. Genes from Supplementary Table 1 (list of all genes in the PRG) not appearing here are not antigen-presenting or cannot be typed for technical reasons (no IMGT exon data available or incomplete resolution of the exon-to-genomic, genomic-to-haplotype alignment steps during PRG construction). *HLA-DRB3* and *HLA-DRB4* copy numbers are variable and linked to DRB1 genotype (neither aspect is modelled by HLA*PRG). Assumedly absent alleles (as determined by linkage with the inferred DRB1 alleles) are shaded in grey, and we note that these carry low quality scores. We detect one trio inconsistency at the HLA-K pseudogene (shaded in bright red), and note that the allele driving the inconsistency carries a low quality score.

## 1 Constructing a PRG for the HLA genes

In this section, we describe how to build a Population Reference Graph (PRG) for the HLA genes (and other genes in IMGT). Basic PRG algorithms and methodology were described in Dilthey, Cox et al. (2015). We briefly recapitulate some important concepts:

- PRGs are derived from multiple sequence alignments (MSAs) of alleles or alternative sequences, such as the 8 extended MHC (xMHC) haplotypes in GRCh38.
- If the number of input sequences is variable across the region to be covered, it is necessary to partition the region into blocks; each individual block contains the same number of sequences across the length of the block. The blocks are processed separately and their graphs later concatenated. For example, for most genes, there are genomic sequences, spanning the entire length of the gene, and additional coding sequences (without intronic / UTR sequences). Each exon therefore becomes a separate block (with an identical number of sequences), as does each intronic region (Fig. 1 b).
- Although not discussed explicitly in Dilthey, Cox et al. (2015), there is no need for the PRG to span a contiguous genomic region; here we build one just from a set of genes, ignoring most of the inter-genic sequence.

### 1.1 Topology of the HLA PRG

The HLA PRG comprises 46 genes, including all classical HLA genes. We have at least one genomic sequence for each gene, and sometimes also coding sequences (see Supplementary Table 1 for a list of genes and input data). For most genes, all available data come from IMGT; however, when there were no IMGT data for a gene (or pseudogene), we have extracted reference sequences directly from the xMHC haplotypes. Input sequences for the HLA PRG are available for download as part of the HLA*PRG data package (http://birch.well.ox.ac.uk/HLA-PRG.tar.gz, list of files: segments.txt).

Sequence data for each gene are transformed into alignment blocks (described below). We connect alignment blocks from different genes with buffer regions, consisting of 2000 bases of undefined sequence (equivalent to ‘N’ characters).

**Figure 1.**
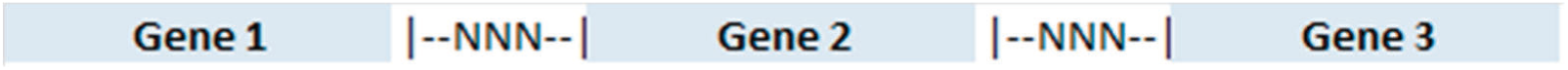
Topology of the HLA PRG: Gene-specific regions are connected with large blocks of undefined sequence (‘N’ characters).

### 1.2 Gene-specific alignment blocks

For each gene, we combine different data sources, when available:

- MSA of genomic sequences (always available), spanning the whole length of the gene.
- MSA of coding sequences, spanning the exons of a gene. Exon boundaries are marked in the alignment.
- xMHC haplotypes from GRCh38, used to embed each gene into a short stretch of genomic context sequence (“padding sequence”; enabling the mapping of reads that span the boundaries of the gene)

For each gene, we harmonize and integrate the available data; and construct a sequence of contiguous multiple sequence alignments.

For each gene, the sequence of MSA blocks comprises (from left to right):

1. Left-padding sequence (xMHC genomic context)
2. Intermittent blocks of intron/UTR and exon sequence, or a monolithic block of genomic sequence.
3. Right-padding sequence (xMHC genomic context)

**Figure 2.**
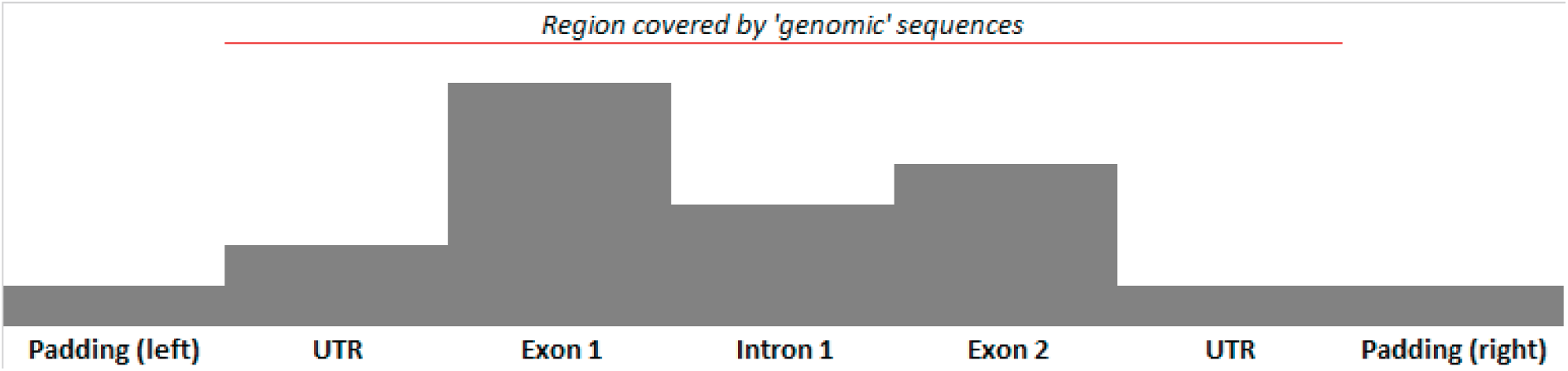
Schematic depiction of the block structure of sequence data for an HLA gene. The Y axis represents the number of sequences going into the corresponding block.

#### 1.2.1 Graphical illustration

Before describing the process for combining and harmonizing gene-specific data in detail, we give a graphical summary and high-level algorithmic of the process and its outcome.

**Figure 3.**
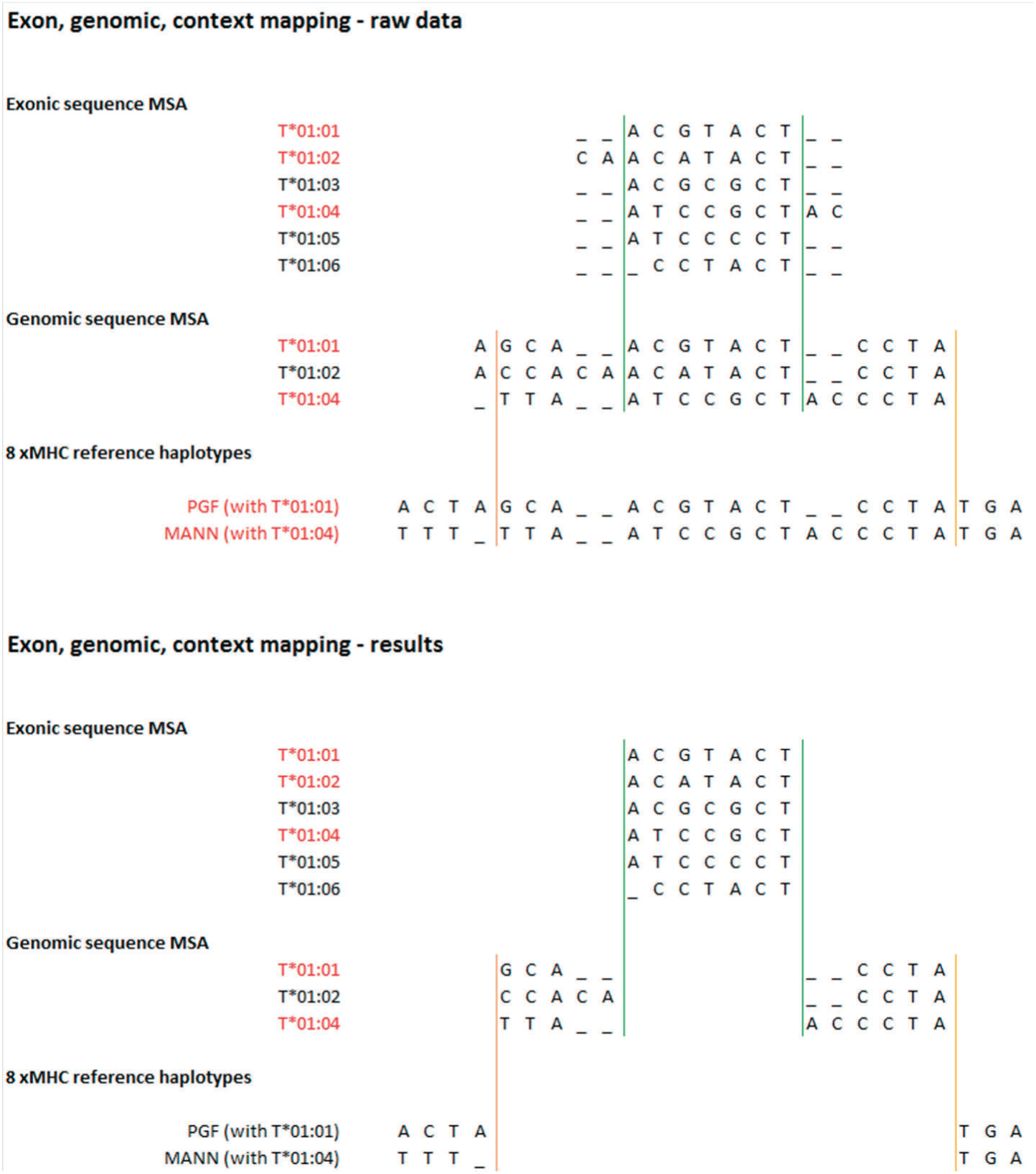
Upper part of figure: input data. Lower part: constructed alignment blocks. We integrate different sources of data based on sequences present at more than one level. Sequence identifiers in red mark the sequences that are present in the next-lower level and that are used to construct a joint coordinate system. The green line marks the boundary of the MSA exon block utilized, the orange line the boundary of the genomic sequence alignment block utilized. The shown sequences are illustrative.

#### 1.2.2 Intuitive description of the MSA merging algorithm

We now give an intuitive description of the MSA merging algorithm.

Each gene is processed independently.

From the description above it is clear that the key challenge is to determine the “switching points” between the different MSAs when constructing the PRG. To give a generic example, we start with the genomic sequences MSA for the gene, and we switch to the exon 1 MSA as soon as possible (because the exon sequences MSA contains more alleles and thus a better representation of sequence diversity). After exon 1 we return to the genomic sequences, and switch to the exon 2 MSA as soon as possible etc. until the whole gene is represented (this example ignores the padding sequences, for which we employ a similar procedure).

The switching points are visualized as vertical lines in Figure 3 of this document (above), and we refer to the contiguous areas of one MSA in which no switch happens as “alignment blocks”. Note that each vertical line has 2 coordinates, one for each MSA that the line connects.

To determine the coordinates of the switching points, we leverage the fact that there are alleles which are represented both in the exon-level MSA and in the genomic-level MSA: if we know that the allele T*1 is represented in both alignments, we should expect to find the exon sequences of T*1 as substrings of the genomic sequence of T*1.

Translating (for each exon independently) the substring match coordinates relative to the un-aligned sequences into alignment space (taking into account the alignments’ gap structure) gives us the <u>alignment</u> coordinates of the switching points between exon and genomic MSAs (and by implication the coordinates of the alignment blocks).

There is additional complexity if there is more than one allele shared between the exonic and genomic MSAs (which is generally the case): there is no guarantee that the switching points computed independently for each shared allele agree. In this case we compute consensus exon alignment blocks: for each exon MSA, we define the left boundary of the corresponding alignment block as the <u>maximum</u> of the left boundaries of all independently computed per-allele alignment blocks (and we proceed analogously for the right boundary). The consensus exon alignment blocks define which areas of the exon MSAs go into the PRG and the corresponding switching points. In a final step, we re-compute MSAs for the genomic sequence alignment blocks for the shared alleles to connect the consensus exon blocks (for each shared allele’s genomic sequence, we extract the corresponding substring in the raw unaligned genomic sequence, and create an MSA of the sequences so-extracted for each block).

#### 1.2.3 Formal description: Base case

We give a formal description for the case of merging the MSA for a single exon into the MSA for surrounding genomic sequence (resulting in 3 alignment blocks). More complex cases follow immediately (see next section). We assume that there are multiple shared alleles (which is almost always the case – the algorithm presented here also works for the one shared allele case).

1. Consensus block coordinates for the exon MSA:

a. Shared alleles: Identify the alleles that are present in both the exon MSA and in the genomic sequence MSA. These alleles will be used for determining the 2 switch points.
b. Initialize *P_L_* = {} and *P_R_* = {}. These two lists will store the coordinates of the allele-specific left and right switch points in the exon MSA (i.e. the entry and exit points in the exon MSA at which the PRG will switch from genomic MSA to exon MSA and back).
c. Allele-specific switch points: For each shared allele, add the exon MSA coordinates of the beginning and the end of the un-aligned allele sequence to *P_L_* and *P_R_*, respectively (for example, if the exon MSA sequence of a shared allele looks like —ACGT…, we add the value 3 to *P_L_* – because there are two gaps in front of the allele exon sequence).
d. Consensus exon alignment block: Set the coordinates of the consensus exon alignment block (in exon MSA space) to *G_L_ =* max(*P_L_*) and *G_R_ =* min(*P_R_*). The consensus exon alignment block so-defined is extracted and goes into the PRG construction process.
2. Genomic MSA alignment blocks:

a. Extract the consensus exon MSA sequence of each shared allele [see Point 1 a) above] – i.e. the allele’s exon sequence bounded by the exon MSA switch points *G_L_* and *G_R_*. Remove all gaps from the sequences so-extracted.
b. Each sequence so-extracted has an exact match in the un-aligned genomic sequence of the corresponding allele.
c. Use the match so-defined to split the un-aligned genomic sequences of the shared alleles. This results in two subsequences per shared allele, corresponding to the left and right genomic MSA alignment blocks.
d. Create the left and right genomic MSA alignment blocks by creating an MSA (e.g. using Muscle (Edgar 2004)) of the left and right split sequences of the shared alleles. The two MSAs so-created go into the PRG construction process.

To obtain a combined PRG, we create 3 PRGs from the 3 MSA blocks (exon consensus, 2 genomic) and merge them accordingly (Dilthey, Cox et al. 2015).

#### 1.2.4 Formal description: Extension cases

<u>Merging genomic into padding sequences:</u> Exactly like the base case.

<u>Merging multiple exons into a genomic sequence MSA:</u> Like the base case, with three modifications:

- We require that the exons be non-overlapping.
- In the base case, we create 2 genomic sequence MSA blocks for the sequences to the left and to the right of the exon. If we have multiple exons, we create additional genomic sequence MSA blocks to cover the space between each pair of subsequent exons (*x* + 1 genomic sequence MSA blocks in total, where *x* is the number of merged exons). The sequences for these genomic MSA blocks are, like in the base case, defined by using the exon consensus block sequences of the shared alleles to split the un-aligned sequences of the shared allele genomic sequences.
- We use an extended definition of “shared alleles”: we additionally require that all members of the “shared alleles” group have the same number of exon sequences that can be mapped uniquely onto their corresponding genomic sequences (this would not be the case, for example, if a particular allele is associated with an exon deletion). If this extended definition is violated, we employ a heuristic that populates the “shared alleles” group in a manner that gives priority to the alleles found on the PGF reference haplotype, and to alleles that have a higher number of exons that can be uniquely mapped.

We use MUSCLE (Edgar 2004) for all MSAs.

## 2 HLA typing

Before describing the algorithms employed in HLA*PRG in detail, we give a high-level summary. HLA typing by HLA*PRG comprises three steps:

1. Read extraction from BAM: Read pairs putatively coming from the HLA genes (i.e. the regions covered by the HLA PRG) are, based on kMer statistics, extracted from an input BAM file.
2. Read-to-graph alignment: Each candidate read is aligned to the HLA PRG. Read mapping quality gives an indication of alignment certainty. If the original alignment from the input BAM file is from a position not covered by the HLA PRG, we re-scale mapping quality accordingly.
3. Inference (6-digit “G” resolution): Only the peptide binding site (PBS) is fully characterized for most alleles present in IMGT/HLA (exons 2 and 3 for class I genes, exon 2 for class II genes). At each locus, we combine all alleles with identical PBS sequences into “clusters” (i.e. one cluster comprises all alleles with identical PBS sequence) and select the most likely pair of allele clusters. This gives HLA types at 6-digit “G” resolution. This process is carried out independently for each locus. Selection of the most likely pair of alleles is based on a likelihood framework.

### 2.1 Step 1: Read extraction from BAM file

We iterate through all read pairs stored in a BAM file and keep only read pairs that

- both reads combined have >30% kMers present in the HLA PRG (positive selection).
- there is at least one read with at least one kMer unique to the HLA PRG, or there is at least one read that has <45% kMers present in regions outside the HLA PRG (negative selection).

For longer reads with lower base qualities, the first criterion is modified to apply to just one read. This feature is activated with the command-line switch `--Hiseq250bp 1`.

To improve performance, we only consider reads

- aligning to chromosome 6, coordinates 28,000,000 – 34,000,000, and unmapped reads
- or chromosome 6 (complete coordinate range) only (`if --Hiseq250bp 1`.).

Thresholds for positive and negative selection are based on exploratory initial experiments. None of the samples used in these experiments are part of the validation cohort.

Positive selection and negative selection are implemented as separate steps – positive selection is slower and requires less memory, whereas negative selection is faster (operating on the results from positive selection only) and memory-intensive (we use parts of the Cortex assembler to compile and hold in memory a list of all kMers present in reference genome regions not covered by the HLA PRG).

For all reads that pass negative and positive selection and that are at least partially aligned (in the BAM file), we score the BAM alignments (separately for each member read of the read pair) corresponding to the likelihood metric presented below, and also store the insert size between the two member pairs. These data will be used later to compare the best alignment from the BAM file with the best HLA PRG alignment.

We use k = 25 for the read extraction step.

### 2.2 Step 2: Read alignment

Alignment of read pairs to the HLA PRG is based on the algorithms presented in (Dilthey, Cox et al. 2015). Let R be the set of all read pairs. To align a read pair r ∈ R consisting of the two member reads r_1_ and r_2_,

1. We find alignments of r_1_ and r_2_ to the graph (independently for r_1_ and r_2_), which we refer to as the sets A(r_1_) and A(r_2_). We refer to these as “member alignments”.

a. The alignment procedure starts by determining strandedness (i.e. determining whether the read is aligned to the + or – strand of the PRG) based on a simple kMer statistic. We compare the number of kMers from the + strand of the read present in the PRG with the number of kMers from the – strand of the read (i.e., the reverse-complemented read sequence) present in the PRG, and assign strandedness accordingly.
b. We identify double-unique kMers (i.e. present once in the read and at only one level in the PRG), which represent exact matches between read and PRG.
c. We proceed by extending each double-unique match with further exact matches to the left and right.
d. The alignment procedure finishes with a local extension step (based on Needleman-Wunsch / Smith-Waterman) that terminates when all bases from the read to be aligned are present in the alignment. Details of a dynamic programming sequence-to-graph alignment algorithm are given in (Dilthey, Cox et al. 2015).
2. We score each element a ∈ ( A(r_1_) ∪ A(r_2_)) according to a simple likelihood function. Alignment a of length L consists of La alignment columns, which we refer to as (c_1_.. c_L_), i.e. 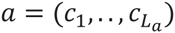 Each such column *c* is an ordered pair (*c_i_*_;1_, *c_i_*_;2_) of two elements, the first of which (*c_i_*_;1_) represents the label and level of the edge in the PRG that the alignment traverses, and the second of which (*c_i_*_;2_) represents the aligned character from the read (both elements can also specify “gap” symbols). We note that *c_i_*_;2_ has an associated base quality score if *c_i_*_;2_ is not the “gap” symbol, and we refer to that quality (after converting the FASTQ Phred score to the probability that the specified base is <u>correct</u>) as q(*c_i_*_;2_). The member alignment score score_member(*a*) is defined as 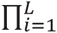 score_pos(*c_i_*), and score_pos(*c_i_*) is defined according to the following table:

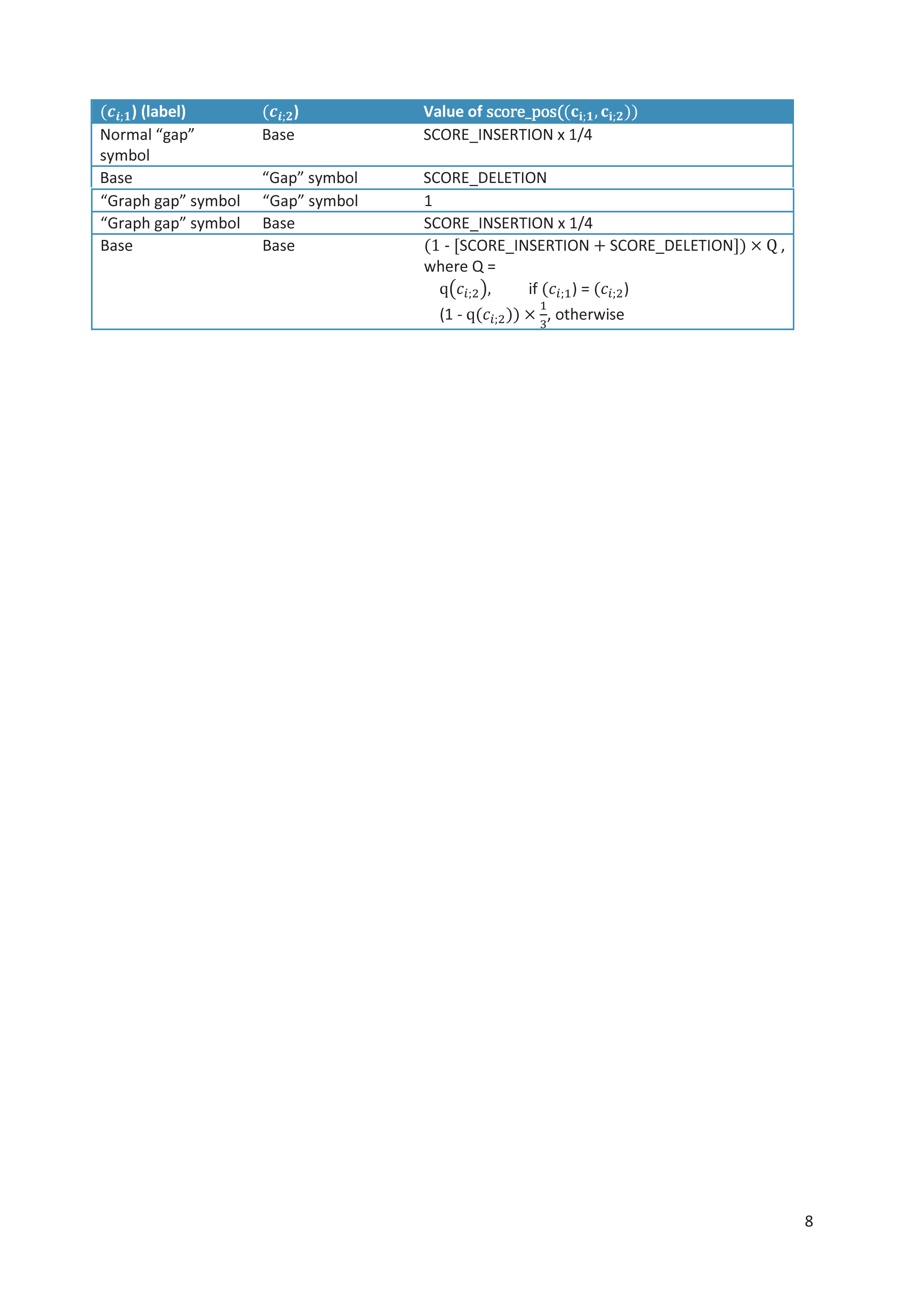
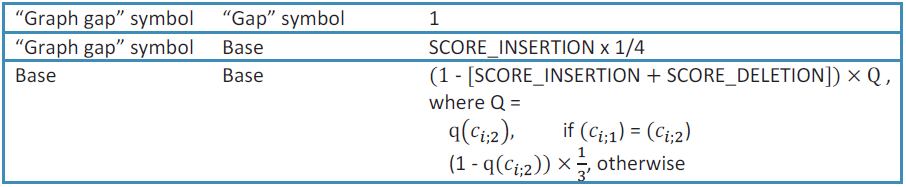 In the current implementation we use SCORE_INSERTION = SCORE_DELETION = 0.01, and we cap 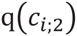 at 0.999.
3. We consider the set 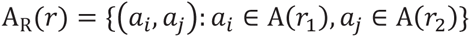 (i.e. the set of all combinations of member read alignments). We refer to each element of A_R_(*r*) as an “alignment” (i.e. the members of A_R_(*r*) are paired alignments). To score a particular combination 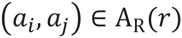, we

a. initialize the score with score_member(*a_i_*) × score_member(*a_j_*).
b. check whether a_i_ and a_j_ are strand-compatible (inverse strands, compatible relative positions of the two member alignments).

i. if not, multiply the score with a penalty factor.
ii. otherwise (i.e. they are strand-compatible), multiply the score with the likelihood of the observed insert size between the two fragments, according to an empirical normal distribution (mean and standard variance of this distribution are heuristically estimated during a preliminary run for each sample).
4. We normalize the scores for all alignments (i.e. the elements of A_R_(*r*)). We refer to the normalized scores as “PRG-only” alignment qualities.
5. As the HLA PRG only covers a fraction of the genome, and as we are operating on a candidate set of reads, we compute a “genomic” alignment quality that takes into account the possibility that the read pair might originate from a region not covered by the HLA PRG. We compute the score for the original alignment as observed in the BAM, unless it is contained (for both member reads) in the region covered by the HLA PRG. Per-read member alignment scores were extracted while filtering the BAM (see Step 1; importantly, the same scoring function is utilized), and mean and standard deviation to calculate the score component for observed insert size based on a normal distribution are now available. We add the score for the original BAM alignment to the list of scores that we normalized during the previous step, and repeat normalization. We call the normalized scores “genomic” alignment qualities (of note, unlike PRG-only alignment qualities, the genomic alignment qualities will typically not sum up to 1 for all possible alignments of a read pair; the original BAM alignment score is part of the normalization procedure, but the alignment itself is not reported). We refer to the genomic mapping quality for an alignment 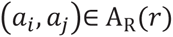 of read *r* as 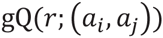.
6. In some cases there is considerable uncertainty in only parts of the alignment (meaning that the alignment score distribution is relatively flat, but that specific combinations of “aligned edge / aligned base” appear in many possible alignments). We initialize an empty hash table and define that, if an element not present in the table is accessed, the element is created and its value is set to 0. We follow the following algorithm:

a. Iterate over all possible alignments (*a_i_*,*a_i_*·) ∈ A_R_(*r*):

i. For a in {*a_i_*,*a_i_*}

1. For all columns in *a*, specifying a traversed edge label and the edge level in the graph, and the aligned sequence character (the character itself and its relative position in the aligned read):

a. Construct a string index containing these properties, as well as a flag specifying whether the alignment *a* is relative to the + or – strand of the PRG.
b. Retrieve the current value of the string index so-constructed from the hash table and increase it by value 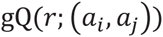.

We can now use the constructed hash table to attach a “per-position” alignment quality value to each column in any of the alignments considered (by constructing the corresponding key for the column under consideration, and retrieving its value).

We use the notation 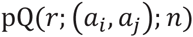 to refer to the per-position alignment quality of column *n* in alignment 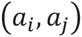 for read *r* (n is a contiguous index over the columns of *a_i_* and *a_j_*, i.e. 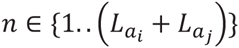.

### 2.3 Step 3: HLA typing

We employ a likelihood framework for HLA tying. That is, we compute the likelihood of the (aligned) read pairs conditional on an assumed underlying pair of HLA types. We consider each locus we want to make an inference for independently.

The inference algorithm (described in this Section) genotypes the HLA genes at the peptide binding site (PBS)-coding positions; for class I genes, these are exons 2 and 3, for class II genes, this is exon 2.

When genotyping at the PBS, we combine (cluster) all reference alleles with identical PBS sequences. No attempt is made to further distinguish between the alleles in a cluster. The results are equivalent to 6-digit “G” resolution HLA typing.

We now give a formal description of the PBS typing process for a specified HLA gene.

#### 2.3.1 Clustering

There is a set of HLA type reference sequences for the gene we are making inference for. These sequences were (by definition) used for the construction of the HLA PRG, i.e. there is an MSA between them; each column of the respective MSA corresponds to one level in the PRG. Also, these sequences are annotated, i.e. intron and exon boundaries (and their positions in the MSA, and hence in the PRG) are available.

We identify the levels of the PRG that correspond to the peptide binding site (PBS) of the gene we are making inference for (these are all levels corresponding to exons 2 and 3 for HLA class I genes, and all levels corresponding to exon 2 for HLA class II genes), and we use the notation PBS_L_ to denote this (ordered) set (each level in the PRG carries a unique identifier, and we require that PBS_L_ is ordered according to order of levels in the PRG; to give a simplified example, PBS_L_ could have the following structure:

{HLA_A_exon_2_pos_l, HLA_A_exon_2_pos_2,…, HLA_A_exon_2_pos_n, HLA_A_exon3_pos_l,…}).

We determine the (ordered) set of unique HLA type reference sequences SEQ^PBS^ across the level set PBS_L_ (i.e. for each HLA type, we extract the nucleotide positions specified by PBS_L_ in their correct order from the corresponding MSAs, concatenate the characters – including gaps, if there are any -, and ensure that the resulting string is present in SEQ_PBS_), and refer to the *i*-th element of SEQ^PBS^ as 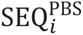. We also refer to these elements in un-concatenated form as “string lists”, in which each PRG level represents one individual member element.

We define 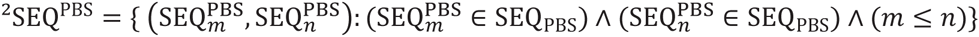 as the set of possible diploid PBS sequences, and use the notation 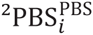 to refer to its *i*–th element.

#### 2.3.2 PBS inference

We now state the inference problem in likelihood terms:

Maximize the likelihood function L_PBS_(*R*| ^2^*S*) over ^2^*S* ∈ ^2^SEQ^PBS^, where *R* is the set of aligned read pairs.

We define

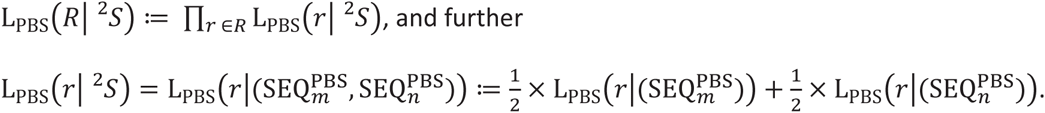

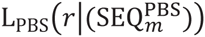 is the likelihood of one read pair, conditional on an assumed underlying (haploid) HLA type sequence (across the PBS).

**Read likelihood: Definition of** 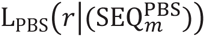

As the PBS sequences we operate on are defined in terms of PRG coordinates, and as the reads we operate on are aligned to the PRG, there is usually a 1:1 correspondence between aligned bases from the read, underlying PRG edges and underlying PRG levels.

However, whenever a read is aligned in a manner that introduces “gap” symbols along the graph dimension, such as for the third position in the following example alignment, this strict correspondence is broken.

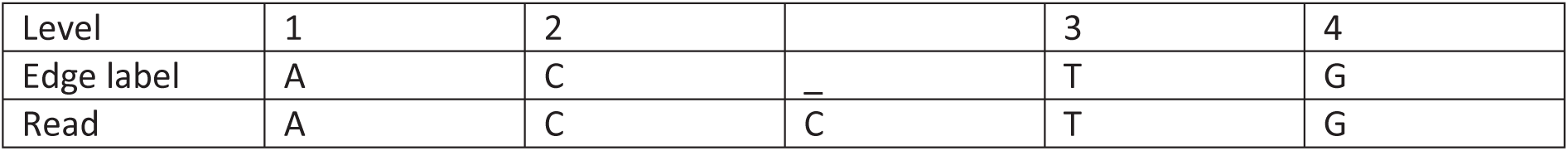

|  |  |  |  |  |  |
| --- | --- | --- | --- | --- | --- |
| Level | 1 | 2 |  | 3 | 4 |
| Edge label | A | C | — | T | G |
| Read | A | C | C | T | G |

We deal with this case by attaching read bases without defined graph level to the preceding alignment column. For the example alignment, we wish to obtain the following result:

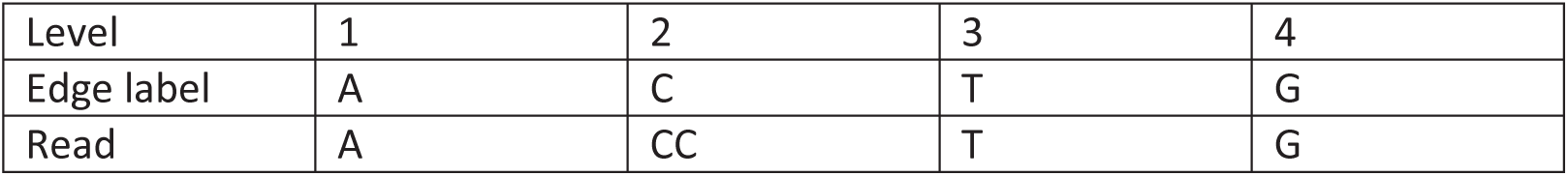

This section describes the technicalities of this transformation and how we proceed to calculate the likelihood.

For the following, note that PRG edges themselves can be labelled with gap characters, such as in the following example:

|  |  |  |  |  |  |
| --- | --- | --- | --- | --- | --- |
| Level | 1 | 2 | 3 | 4 | 5 |
| Edge label | T | A | — | T | G |
| Read | T | A | — | T | G |

We refer to these gap-labelled edges as “graph gaps” and note that “graph gaps” will not be transformed during the following steps.

1. Best alignment: To define 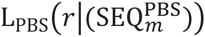, we select the optimal quality alignment (*a_i_*, *a_j_*·) from A_R_(*r*) (i.e. the alignment achieving the maximum 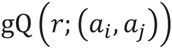 value).
2. Per-read alignment concatenation: We concatenate the two per-read alignments (*a_i_*, *a_j_*) to obtain a combined alignment vector: We define the combined column set *C* as the concatenation of the columns of *a_i_* and *a_j_* (i.e. 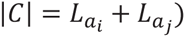). From the definitions given earlier, we recapitulate that each element *c_i_* ∈ *C* consists of two sub-elements, i.e. *c_i_* = (*c_i_*_;1_, *c_i_*_;2_). The first element *c_i_*_;1_ specifies label and level of the edge traversed in the PRG, and the second element *c_i_*_;2_ specifies the corresponding base from the aligned read (both sub-elements can also specify “gaps”).
3. Removal of “gaps”: We create the set *C* by removing all columns from *C* that specify “gap” symbols along the graph dimension (but not proper “graph gaps”), using the following algorithm:

a. Set *C*′ = *C*
b. If there is no element (*c_i_*_;1_, *c_i_*_;2_) ∈ *C*′ that does *not* specify a gap for *c_i_*_;1_, set *C*′ = {} and return *C*′ (note that we differentiate between “gap” symbols and “graph gap” symbols, the latter of which indicate traversed edges in the graph that are themselves labelled with “graph gap” symbols; we only want to remove proper “gap” columns).
c. Traverse *C*′ from left to right and find the first column *c_i_* with (*c_i_*_;1_, *c_i_*_;2_) specifying a “gap” for *c_i_*_;1_.

i. If *i* > 1 and the column *c_i–_*_1_ = (*c_i–_*_1;1_, *c_i–_*_1;2_) to the left of the current position does not specify a graph “gap”, attach *c_i_*_;2_ to *c_i–_*_1;2_ and remove *c_i_* from *C*′. Go back to Step *c*.
ii. If I = 1, attach *c_i_*_;2_ to the beginning of *c_i_*_+1;2_, remove c from *C*′, and go back to Step *c*.
d. If no such columns were found during the last iteration of Step *c*, terminate and return *C*′. We note that *C* may now contain elements (*c_i_*_;1_, *c_i_*_;2_) with sub-elements *c_i_*_;2_ longer than one character.
4. If *C*′ = {}, set 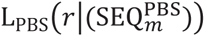 to 1 and return.
5. Each remaining column in *C*′ has one and only one corresponding level in the PRG. We remove all columns with levels not present in PBS_L_. If *C*′ is now the empty set, set 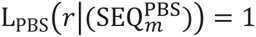 and return.
6. We define
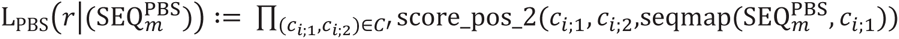, where

a. score_pos_2 is defined below
b. seqmap is a function that returns the underlying genotype of HLA type sequence 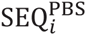 at the PRG level specified by the level component of *c_i_*_;1_.

| GT | $(c_{i;2})$ | Value of $\text{score\_pos\_2}((c_{i;1}, c_{i;2}), \text{GT})$ |
| --- | --- | --- |
| Gap | (Non-gap) base(s) | $(\text{SCORE\_INSERTION} \times \frac{1}{4})^{\text{LENGTH}((c_{i;2}))}$ |
| Gap | “Gap” symbol | 1 |
| Base | String | Algorithm:<br>forReturn := 1<br>**IF** SUBSTRING($(c_{i;2}), 0, 1$) == “_”<br>forReturn *= SCORE_DELETION<br>**ELSE**<br>forReturn *=<br>$(1 - [\text{SCORE\_INSERTION} + \text{SCORE\_DELETION}]) \times Q$, |

#### 2.3.3 Posterior probabilities and best-guess extraction

We normalize 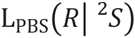 to obtain posterior probabilities over possible diploid genotypes. To obtain a “best guess” of two individual alleles, we employ the same procedure as for HLA*IMP:02 (Dilthey, Leslie et al. 2013). Briefly, for each allele in ^2^*S*, we first compute the probability of occurring at least once (integrating over all allele pair probabilities). We fix the maximum allele as the first best-guess allele, and use this marginal probability as our quality score for allele 1. Having fixed the first allele, we now extract all pairs that contain at least one instance of allele 1, and select the pair with the maximum absolute posterior probability. We use the second allele as the second best-guess allele, and use the absolute probability of the allele pair as the quality score for allele 2.

### 2.4 Runtime and computational resources

We measured runtime and RAM usage for all steps of the HLA typing process for NA12878 (Platinum and 1000 Genomes):

#### 2.4.1 NA12878 Platinum 55× 2 × 100bp

2.4.1 NA12878 Platinum 55x 2 x 100bp
| Step |  | Clock (wall) time | CPU time (user time, all threads combined) | Maximum RAM | Threads | Comments |
| --- | --- | --- | --- | --- | --- | --- |
| 1.1 | Positive filtering | 7h | 6.9h | 12G | 1 |  |
| 1.2 | Negative filtering | 29m | 0.35h | 75GB | 1 |  |
| 2 | Read alignment | 3h | 32.5h | 70G | 40 |  |
| 3 | HLA type inference | 9m | 0.46h | 1.6G | 32 |  |

#### 2.4.2 NA12878 1000 Genomes 63× 2 × 250bp

2.4.2 NA12878 1000 Genomes 63x 2 x 250bp
| Step |  | Clock (wall) time | CPU time (user time, all threads combined) | Maximum RAM | Threads | Comments |
| --- | --- | --- | --- | --- | --- | --- |
| 1.1 | Positive filtering | 10.5h | 10.4h | 13G | 1 | Run with<br>--HiSeq250bp<br>1 |
| 1.2 | Negative filtering | 39m | 0.5h | 75GB | 1 |  |
| 2 | Read alignment | 6h | 215h | 70GB | 40 |  |
| 3 | HLA type inference | 8m | 0.3h | 1.4G | 32 |  |

## 3 Validation

Due to the complexities of HLA type nomenclature, validating and comparing the performance of different HLA type inference algorithms in a fair manner is not always straightforward. Below we describe our validation approach in detail.

### 3.1 Input data format

We use BAM files, the current standard for storing genomic data, as the starting point for all analyses and comparisons between algorithms.

If necessary, we create BAMs from raw FASTQ data with BWA 0.6.2. For the 1000 Genomes samples in particular, we use BAM files downloaded from the 1000 Genomes website (see Section “Data” for the URLs).

The first step of the HLAPRG pipeline, positive selection, operates on BAM files. HLAreporter and PHLAT require FASTQ input, which we extract, using Picard (http://picard.sourceforge.net), from the BAM files analyzed by HLA*PRG.

### 3.2 4-digit, 6-digit and ambiguous HLA types

Performance assessment and performance comparisons are complicated by the fact that there are multiple resolutions for HLA types:

- 6-digit HLA types (“high resolution”) specify the sequence of all exons of the HLA gene.
- 4-digit HLA types (“intermediary resolution”) specify the primary structure of the HLA protein, i.e. they specify the amino acids encoded by the exons of the HLA gene.
- 6-digit “G” types specify the sequence of the exons encoding the peptide binding site (PBS) region of the HLA gene (exons 2 and 3 for HLA class I genes and exon 2 for HLA class II genes). The reference list of 6-digit “G” groups is available at the IMGT/HLA website: http://hla.alleles.org/wmda/hlanomg.txt.
- 2-and 8-digit HLA types are not relevant in the context of this publication,

The current gold-standard for HLA typing is sequence-based typing (SBT), and most of the validation data used here was generated by SBT. SBT typically gives results at 6-digit “G” resolution. We briefly highlight some important properties of 6-digit “G” codes:

- A 6-digit “G” code is often ambiguous: that is, many individual 6-digit “G” codes map to a list of possible underlying 6-digit (non-G) codes (which are differentiated by polymorphism in non-PBS exons – see the list provided by IMGT/HLA).
- A 6-digit “G” code can map to multiple 4-digit HLA codes (if a polymorphism in non-PBS exons Leads to an amino acid change).

### 3.3 Ambiguity in the validation data

When dealing with ambiguous validation data (which is almost always the case, see above), we explicitly carry the ambiguity through the validation process and validate at the highest resolution specified by the validation data.

The following table lists some examples (a formal definition is given below):

| Validation Allele 1<br>(possibly a set of alleles) | Validation Allele 2<br>(possibly a set of alleles) | Inferred Allele 1 | Inferred Allele 2 | Number of inferred alleles counted as correct |
| --- | --- | --- | --- | --- |
| 02:01:01;02:01:02 | 01:01:01 | 01:01:01 | 02:01:01 | 2 |
| 02:01:01;02:01:02 | 01:01:01 | 01:01:01 | 02:01:03 | 1 |
| 02:01 | 01:01:01 | 01:01:01 | 02:01:03 | 2 |

### 3.4 Ambiguity in HLA*PRG results (and these of other algorithms)

Similar to SBT, the primary output from HLA*PRG is at 6-digit “G” resolution.

That is, ambiguity exists not only in the validation data, but also in the inference dataset, but (at least for HLA*PRG and SBT validation data) the allele groups found in the inference and validation data will be identical.

This, however, is not necessarily the case for the other programs we benchmark HLA*PRG against. We generally preserve ambiguity in the inference results as specified by the other programs, and we count an inferred allele (or ambiguous allele group) as correct if and only if one of the contained alleles validates successfully against one of the specified validation alleles (or ambiguous validation allele groups). Importantly, an inferred allele that is at a lower resolution than the validation allele will never validate successfully (because the validation data determines validation resolution; but see below for a 2^nd^ validation metric).

We give some examples for ambiguity and different HLA type resolutions in the inference set:

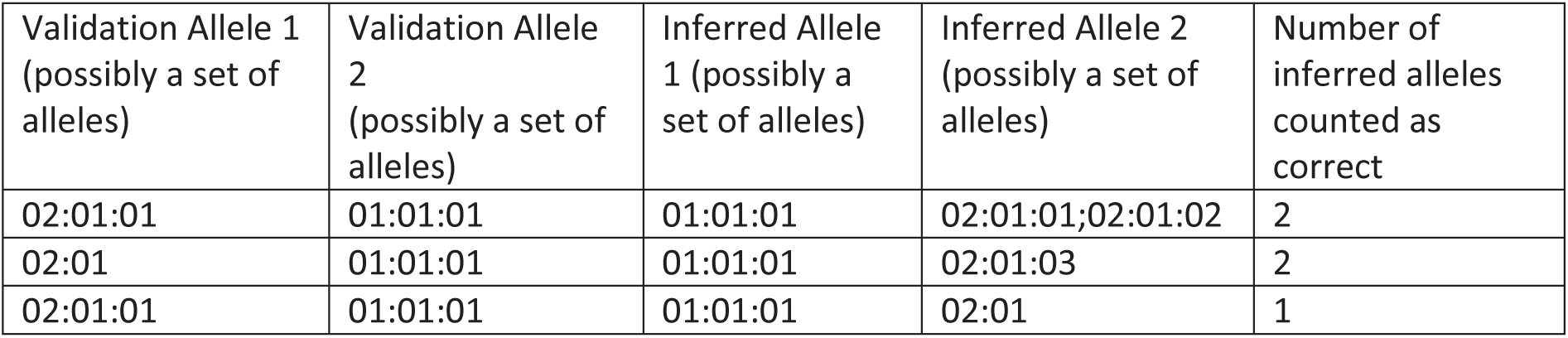

#### 3.4.1 “4-digit” validation

The approach described above is arguably biased against programs that output 4-digit alleles instead of 6-digit allele groups if resolution of ambiguity is not possible (both points apply to PHLAT).

Therefore we also consider an additional metric of accuracy for which we reduce all validation alleles (and by implication all inferred alleles) to 4-digit resolution.

We note that this reduction, by definition, doesn’t result in proper 4-digit alleles: as most of the original validation data is at 6-digit “G” resolution, amino acid positions outside exons 2 and 3 (or 2 for class II) remain undetermined. After applying our reduction, an inferred allele will only validate successfully if it has the same amino acid sequence over exons 2 and 3 (or 2 for class II) as the validation allele.

### 3.5 Formal description

We now give an algorithmic description of our validation approach. The “4-digit evaluation” described in the previous paragraph follows immediately by converting all validation alleles to 4-digit accuracy.

*HLA-X data for individual Y:*

|  | Allele 1 | Allele 2 |
| --- | --- | --- |
| Inferred | $I_1 = \{\text{allele1}, \text{allele2}, \dots\}$ | $I_2 = \{\text{allele1}, \text{allele2}, \dots\}$ |
| Validation | $V_1 = \{\text{allele1}, \text{allele2}, \dots\}$ | $V_2 = \{\text{allele1}, \text{allele2}, \dots\}$ |

We note again that *I*_1_, *I*_2_, *V*_1_, *V*_2_ are groups of alleles that can, without loss of generality, consist of only one member.

We now define the number of correctly inferred alleles as

correct (*I*_1_, *I*_2_, *V*_1_, *V*_2_): = max(correct2(*I*_1_, *I*_2_, *V*_1_, *V*_2_) correct2(*I*_1_, *I*_2_, *V*_2_, *V*_1_)),

and we define

correct2(*I_x_*, *I_y_*, *V_x_*, *V_y_*): = correct1(*I_x_*, *V_x_*) + correctl(*I_y_*,*V_y_*),

and finally

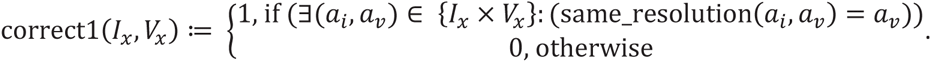

same_resolution(*a_i_*, *a_v_*) is a function that transforms (and then returns) *a_i_* to the same resolution as *a_v_* (either by removing digit groups or by adding “:00” groups). We give three examples:

| $a_i$ | $a_v$ | $\text{same\_resolution}(a_i, a_v)$ |
| --- | --- | --- |
| 01:02:01 | 03:01:04 | 01:02:01 |
| 01:02 | 03:01:04 | 01:02:00 |
| 01:02:01 | 03:01 | 01:02 |

We also integrate the notion of “missingness”. “Missingness” can arise, for example, due to only one validation allele (group) being specified for a sample, or due to the removal of one inferred allele (group) because of the application of a posterior probability call-threshold which the removed allele (group) doesn’t meet.

We define correct1(*I_x_*,*V_x_*) as 0 for all instances in which *I_x_* or *V_y_* are set to “missing”.

The number of “called” alleles (always 2 when there is no missingness) is defined as

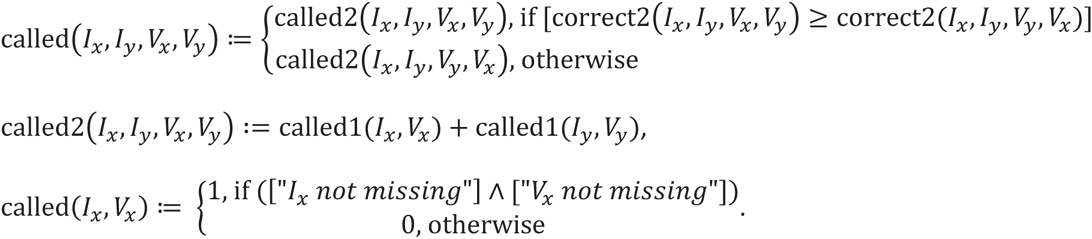

Accuracy and call rate for a set of samples are computed by summing over the number of correct alleles: accuracy is computed by dividing the sum of correct alleles by the sum of called alleles, and call rate is computed by dividing the number of called alleles by [“number of samples” x 2]

### 3.6 PHLAT-specific details

The output from PHLAT is validated as it is produced. To account for the fact that PHLAT emits lower-resolution alleles in cases of ambiguity, we report the “4-digit” validation metric. During the 1000 Genomes validation experiment, PHLAT consistently failed to produce output for one sample, which we count as “not called”.

### 3.7 HLAreporter-specific details

HLAreporter sometimes emits alleles specified at 6-digit “G” resolution. We transform these into the corresponding 6-digit allele groups as specified by IMGT (http://hla.alleles.org/wmda/hlanomg.txt).

We observe that HLAreporter often generates empty call files. These are interpreted as “missing”, will lower the measured call rate and will not contribute to accuracy metrics.

The authors of HLAreporter suggested modifications to deal with 2 × 250bp reads. Specifically, they recommended generating a new set of pseudo-reads by splitting each 250bp read in half. To give an example, the read pair (original_read_1, original_read_2) would generate the two new pseudo read pairs: (firstHalf_original_read_1, secondHalf_original_read_1), (firstHalf_original_read_2, second_half_original_read_2). Results without this modification were very similar.

HLAreporter call files are processed according to the following algorithm:

- For class I loci, we search for the string “Allele pair”, and extract the next two lines. If the first line specifies a valid allele (with the right locus identifier), we use the specified allele as allele 1 – otherwise we specify allele 1 = allele 2 = “missing”. If the second line also specifies a valid allele, we use the specified allele as allele 2. Finally, if allele 1 was set but not allele 2, we define allele 2 = “missing”.
- For class II loci, we search for the occurrence of the string “Allele” at the beginning of a line, and read the subsequent lines (which specify alleles) until we hit another “Allele” string or the string “HLA data quality profile” (which marks the end of the allele list). We define allele 1 as the list of all alleles specified after “Allele” (i.e. allele 1 is potentially a list of alleles). If we find another “Allele” string, we repeat the same process, but now use the found alleles to define allele 2. Finally, if allele 1 was set but not allele 2, we define allele 2 = “missing”.

If the call file contains a “low data quality” warning (as it almost always does in our case), we assign low posterior probabilities to the extracted alleles.

### 3.8 DRB1 correction for NA19238, NA19239

#### 3.8.1 NA19238

The following *HLA-DRB1* types are specified in the original (non-corrected) validation data:

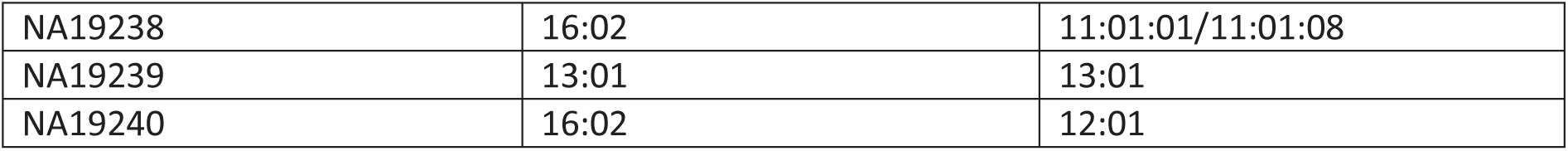

NA19240 is the child of NA19238 and NA19239. We noted that these HLA types are transmission-incompatible. The most likely scenario is that either one of the 13:01 alleles of NA19239 is an error, or the 12:01 allele of NA19240.

HLA*PRG also infers that NA19239 has one 12:01 allele, which would be transmission-consistent.

High-resolution sequence-based re-typing confirms that the correct DRB1 genotype is

|  |  |  |
| --- | --- | --- |
| NA19239 | 12:01:01;12:10 | 13:01:01 |

#### 3.8.2 NA19239

For NA19238, the originally specified genotype for *HLA-DRB1* was 16:02 / 11:01:01;11:01:08.

HLA*PRG predicts 11:01:02 / 16:23.

High-resolution sequence-based typing confirms that the *DRB1* genotype for this sample is 11:01:02 / 16:02:01, i.e. one of the two discrepancies for NA19239 between HLA*PRG and the original reference data was driven by an error in the original reference data.

## 4 Data

### 4.1 HLA types

HLA types for 1000 Genomes Samples (Gourraud, Khankhanian et al. 2014) were downloaded from ftp://ftp.1000genomes.ebi.ac.uk/vol1/ftp/technical/working/20140725_hla_genotypes/.

High-resolution HLA typing for some samples present in the HapMap cohort was available from Dilthey, Leslie et al. (2013).

Two *HLA-B* alleles for NA12891 were added from Erlich, Jia et al. (2011).

We identified two errors in the 1000 Genomes Data and changed the allele accordingly (see Section 3.8).

#### 4.1.1 Validated HLA types for Platinum samples

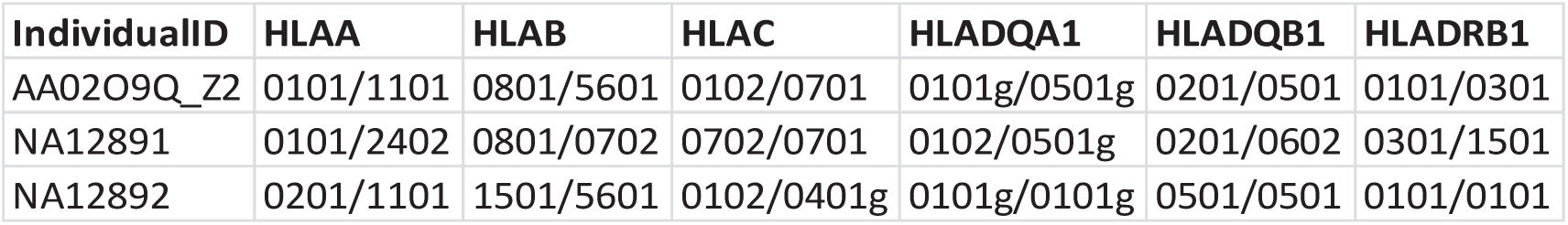

#### 4.1.2 Validated HLA types for 1000 Genomes samples

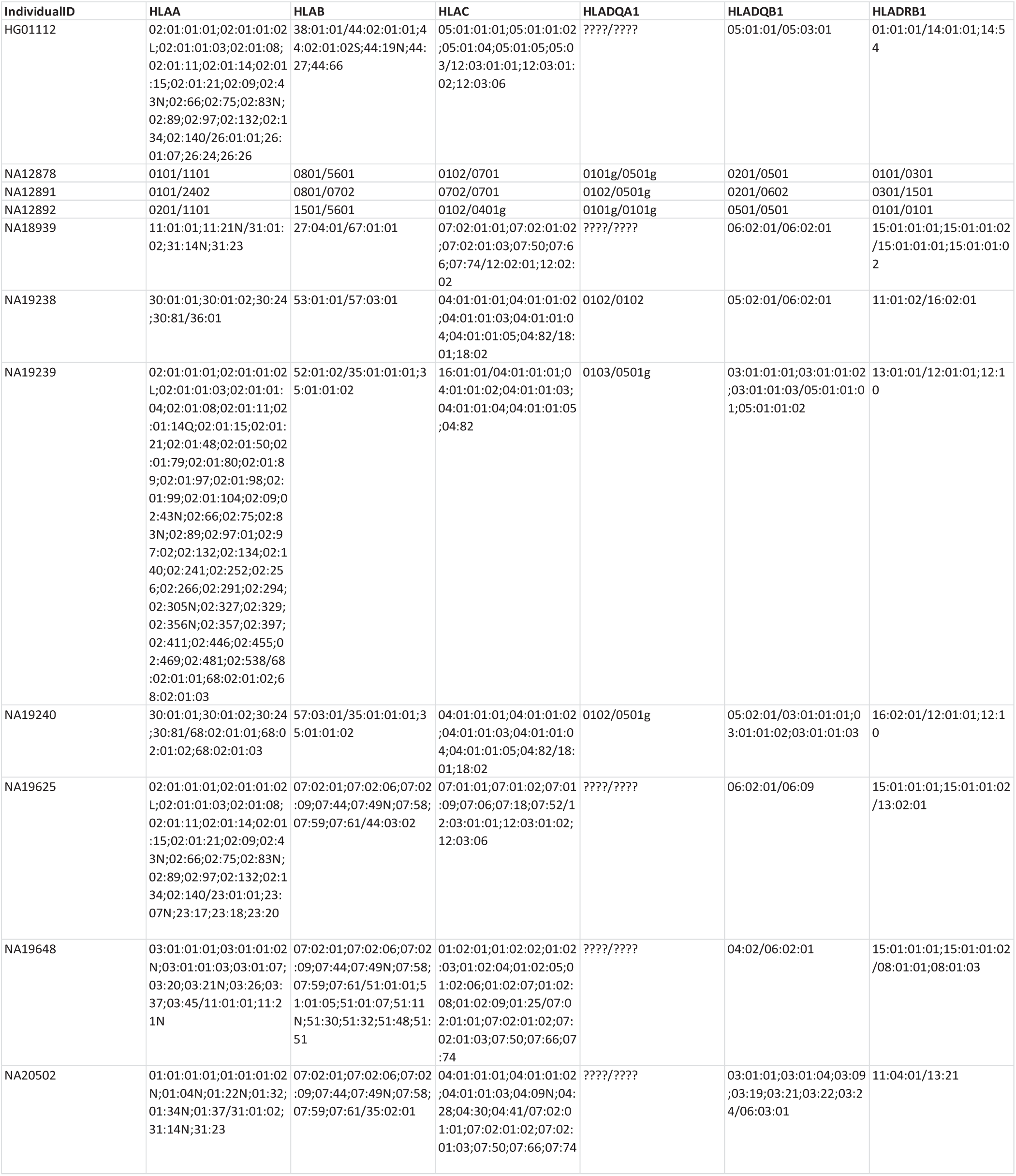

#### 4.1.3 Validated HLA types for HapMap exome samples

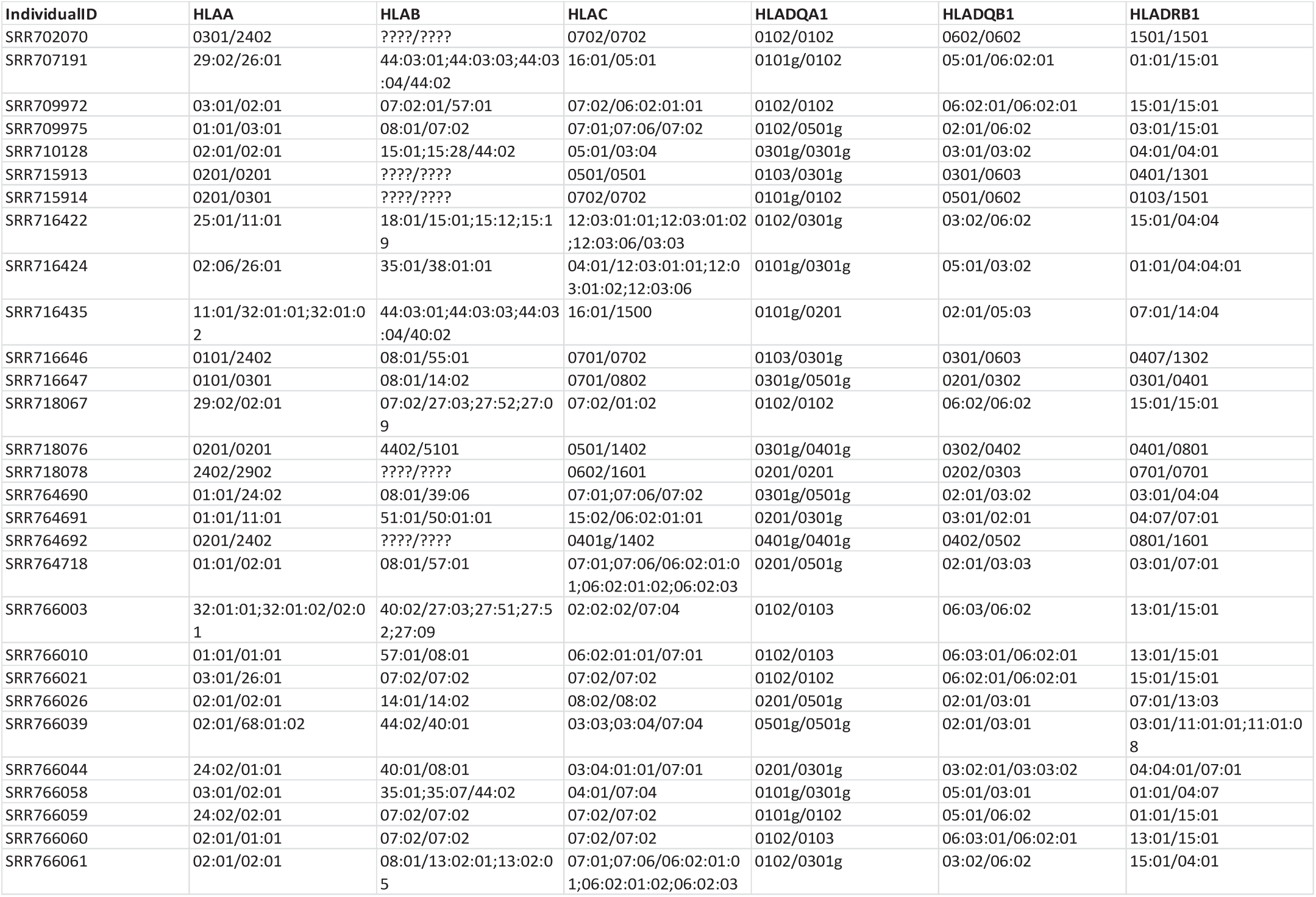

| IndividualID | HLA A | HLA B | HLA C | HLA DQA1 | HLA DQB1 | HLA DRB1 |
| --- | --- | --- | --- | --- | --- | --- |
| SRR702070 | 0301/2402 | ????/???? | 0702/0702 | 0102/0102 | 0602/0602 | 1501/1501 |
| SRR707191 | 29:02/26:01 | 44:03:01;44:03:03;44:03:04/44:02 | 16:01/05:01 | 0101g/0102 | 05:01/06:02:01 | 01:01/15:01 |
| SRR709972 | 03:01/02:01 | 07:02:01/57:01 | 07:02/06:02:01:01 | 0102/0102 | 06:02:01/06:02:01 | 15:01/15:01 |
| SRR709975 | 01:01/03:01 | 08:01/07:02 | 07:01;07:06/07:02 | 0102/0501g | 02:01/06:02 | 03:01/15:01 |
| SRR710128 | 02:01/02:01 | 15:01;15:28/44:02 | 05:01/03:04 | 0301g/0301g | 03:01/03:02 | 04:01/04:01 |
| SRR715913 | 0201/0201 | ????/???? | 0501/0501 | 0103/0301g | 0301/0603 | 0401/1301 |
| SRR715914 | 0201/0301 | ????/???? | 0702/0702 | 0101g/0102 | 0501/0602 | 0103/1501 |
| SRR716422 | 25:01/11:01 | 18:01/15:01;15:12;15:19 | 12:03:01:01;12:03:01:02;12:03:06/03:03 | 0102/0301g | 03:02/06:02 | 15:01/04:04 |
| SRR716424 | 02:06/26:01 | 35:01/38:01:01 | 04:01/12:03:01:01;12:03:01:02;12:03:06 | 0101g/0301g | 05:01/03:02 | 01:01/04:04:01 |
| SRR716435 | 11:01/32:01:01;32:01:02 | 44:03:01;44:03:03;44:03:04/40:02 | 16:01/1500 | 0101g/0201 | 02:01/05:03 | 07:01/14:04 |
| SRR716646 | 0101/2402 | 08:01/55:01 | 0701/0702 | 0103/0301g | 0301/0603 | 0407/1302 |
| SRR716647 | 0101/0301 | 08:01/14:02 | 0701/0802 | 0301g/0501g | 0201/0302 | 0301/0401 |
| SRR718067 | 29:02/02:01 | 07:02/27:03;27:52;27:09 | 07:02/01:02 | 0102/0102 | 06:02/06:02 | 15:01/15:01 |
| SRR718076 | 0201/0201 | 4402/5101 | 0501/1402 | 0301g/0401g | 0302/0402 | 0401/0801 |
| SRR718078 | 2402/2902 | ????/???? | 0602/1601 | 0201/0201 | 0202/0303 | 0701/0701 |
| SRR764690 | 01:01/24:02 | 08:01/39:06 | 07:01;07:06/07:02 | 0301g/0501g | 02:01/03:02 | 03:01/04:04 |
| SRR764691 | 01:01/11:01 | 51:01/50:01:01 | 15:02/06:02:01:01 | 0201/0301g | 03:01/02:01 | 04:07/07:01 |
| SRR764692 | 0201/2402 | ????/???? | 0401g/1402 | 0401g/0401g | 0402/0502 | 0801/1601 |
| SRR764718 | 01:01/02:01 | 08:01/57:01 | 07:01;07:06/06:02:01:01;06:02:01:02;06:02:03 | 0201/0501g | 02:01/03:03 | 03:01/07:01 |
| SRR766003 | 32:01:01;32:01:02/02:01 | 40:02/27:03;27:51;27:52;27:09 | 02:02:02/07:04 | 0102/0103 | 06:03/06:02 | 13:01/15:01 |
| SRR766010 | 01:01/01:01 | 57:01/08:01 | 06:02:01:01/07:01 | 0102/0103 | 06:03:01/06:02:01 | 13:01/15:01 |
| SRR766021 | 03:01/26:01 | 07:02/07:02 | 07:02/07:02 | 0102/0102 | 06:02:01/06:02:01 | 15:01/15:01 |
| SRR766026 | 02:01/02:01 | 14:01/14:02 | 08:02/08:02 | 0201/0501g | 02:01/03:01 | 07:01/13:03 |
| SRR766039 | 02:01/68:01:02 | 44:02/40:01 | 03:03;03:04/07:04 | 0501g/0501g | 02:01/03:01 | 03:01/11:01:01;11:01:08 |
| SRR766044 | 24:02/01:01 | 40:01/08:01 | 03:04:01:01/07:01 | 0201/0301g | 03:02:01/03:03:02 | 04:04:01/07:01 |
| SRR766058 | 03:01/02:01 | 35:01;35:07/44:02 | 04:01/07:04 | 0101g/0301g | 05:01/03:01 | 01:01/04:07 |
| SRR766059 | 24:02/02:01 | 07:02/07:02 | 07:02/07:02 | 0101g/0102 | 05:01/06:02 | 01:01/15:01 |
| SRR766060 | 02:01/01:01 | 07:02/07:02 | 07:02/07:02 | 0102/0103 | 06:03:01/06:02:01 | 13:01/15:01 |
| SRR766061 | 02:01/02:01 | 08:01/13:02:01;13:02:05 | 07:01;07:06/06:02:01:01;06:02:01:02;06:02:03 | 0102/0301g | 03:02/06:02 | 15:01/04:01 |

### 4.2 Next-generation sequencing data

#### 4.2.1 NA12878, NA12891, NA12892 Platinum

Read data for NA12878, NA12891 and NA12892 from the Illumina Platinum

(http://www.illumina.com/platinumgenomes/) genomes project (HiSeq 2000, ~60× coverage, 100-bp paired-end reads) were obtained from the European Bioinformatics Institute (http://www.ebi.ac.uk/ena/data/view/ERP001960).

#### 4.2.2 1000 Genomes High-Coverage

BAM files were downloaded from

ftp://ftp.1000genomes.ebi.ac.uk/vol1/ftp/phase3/data/teSAMPLE_ID}/high_coverage_alignment/ for the following sample IDs:

HG00096

HG03642

HG00268

HG03742

HG00419

NA12878

HG00759

NA12891

HG01051

NA12892

HG01112

NA18525

HG01500

NA18939

HG01565

NA19017

HG01583

NA19238

HG01595

NA19239

HG01879

NA19240

HG02568

NA19625

HG02922

NA19648

HG03006

NA20502

HG03052

NA20845

#### 4.2.3 HapMap Exome Data

FASTQ files for HapMap samples were downloaded from the Sequence Read Archive:

SRR701474 NA11992

SRR702070 NA12873

SRR715913 NA12812

SRR716435 NA12234

SRR718077 NA12813

SRR764691 NA12156

SRR766010 NA11995

SRR766058 NA12144

SRR701475 NA11994

SRR707191 NA11993

SRR715914 NA12814

SRR716646 NA12815

SRR718078 NA12813

SRR764692 NA12874

SRR766021 NA11881

SRR766059 NA12004

SRR702067 NA12154

SRR709972 NA06985

SRR716422 NA12006

SRR716647 NA12872

SRR742200 NA12046E

SRR764693 NA12414

SRR766026 NA11830

SRR766060 NA12044

SRR702068 NA12155

SRR709975 NA11831

SRR716423 NA12043

SRR718067 NA12005

SRR764689 NA07357

SRR764718 NA07056

SRR766039 NA07000

SRR766061 NA12003

SRR702069 NA12489

SRR710128 NA11829

SRR716424 NA12043

SRR718076 NA12762

SRR764690 NA07357

SRR766003 NA11832

SRR766044 NA10851

SRR767596 NA12046E

